# Comparative transcriptomics and metabolomics reveal specialized metabolite drought stress responses in switchgrass (*Panicum virgatum* L.)

**DOI:** 10.1101/2022.04.20.488672

**Authors:** Kira Tiedge, Xingxing Li, Amy T. Merrill, Danielle Davisson, Yuxuan Chen, Ping Yu, Dean J. Tantillo, Robert L. Last, Philipp Zerbe

## Abstract

Switchgrass (*Panicum virgatum*) is a bioenergy model crop valued for its energy efficiency and drought tolerance resilience. The related monocot species rice (*Oryza sativa*) and maize (*Zea mays*) deploy species-specific, specialized metabolites as core stress defenses. By contrast, specialized chemical defenses in switchgrass are largely unknown.
To investigate specialized metabolic drought responses in switchgrass, we integrated tissue-specific transcriptome and metabolite analyses of the genotypes Alamo and Cave-in-Rock that feature different drought tolerance.
The more drought-susceptible Cave-in-Rock featured an earlier onset of transcriptomic changes and significantly more differentially expressed genes in response to drought compared to Alamo. Specialized pathways showed moderate differential expression compared to pronounced transcriptomic alterations in carbohydrate and amino acid metabolism. However, diterpenoid-biosynthetic genes showed drought-inducible expression in Alamo roots, contrasting largely unaltered triterpenoid and phenylpropanoid pathways. Metabolomic analyses identified common and genotype-specific flavonoids and terpenoids. Consistent with transcriptomic alterations, several root diterpenoids showed significant drought-induced accumulation, whereas triterpenoid abundance remained predominantly unchanged. Structural analysis of drought-responsive root diterpenoids verified these metabolites as oxygenated furanoditerpenoids.
Drought-dependent transcriptome and metabolite profiles provide the foundation to understand the molecular mechanisms underlying switchgrass environmental resilience. Accumulation of specialized root diterpenoids and corresponding pathway transcripts supports a role in drought stress tolerance for these compounds.

**Significance statement:** With an increasing demand for renewable energy opposed by rising climate-driven crop losses, understanding, and leveraging plant natural defenses can enable the development of sustainable crop production systems. Here, we integrated comparative transcriptomics and metabolomics analyses to gain a detailed understanding of the diversity and physiological relevance of specialized metabolites in upland and lowland switchgrass ecotypes and provide resources for future investigations of drought response mechanisms in switchgrass.

## Introduction

Water scarcity exacerbated by climate change threatens biofuel and food crop production across the world (Pokhrel et al. 2021; Challinor et al. 2014; Kim et al. 2019b). In the U.S., about one-third of all counties are currently designated as crop loss disaster areas through drought by the U.S. Department of Agriculture (USDA Farm Service Agency, 2021). Crop production is further impacted by climate-associated increases in pest and pathogen damage (Newbery et al. 2016), calling for new solutions to develop crops that can withstand current and future climate conditions.

The perennial grass switchgrass (*Panicum virgatum*) is a characteristic species of North American tallgrass prairie land and of agroeconomic value as a C_4_ lignocellulosic feedstock (McLaughlin et al. 1999). A high net-energy yield and environmental resilience make switchgrass economically viable for biofuel production on marginal lands with reduced agricultural inputs. Two major switchgrass ecotypes, Northern upland and Southern lowland, differ in climatic and geographical adaptation, morphological characteristics and genetic architecture (Ayyappan et al. 2017; Lowry et al. 2014). Upland ecotypes are mostly octoploid (2n=8x=72), whereas lowland ecotypes are predominantly tetraploid (2n=4x=36) and feature taller phenotypes with thicker stems and a later flowering time (Casler et al. 2011). The recent development of genome resources for the allotetraploid lowland ecotype Alamo (∼1.23 Gb, NCBI:txid38727) (Lovell et al. 2021) now provides the foundation needed to investigate genetic and biochemical mechanisms underlying switchgrass environmental resilience. Indeed, genomic analysis of 732 switchgrass genotypes across 1,800 km latitude range revealed an extensive correlation of genomic architecture to climatic adaptation (Lovell et al. 2021). Comparative morphological and physiological analysis of 49 upland and lowland ecotypes showed significant differences in the drought tolerance of different switchgrass ecotypes (Liu et al. 2015). Large-scale transcriptomic changes were also observed, including a drought-induced down-regulation of photosynthetic genes, consistent with physiological responses such as reduced leaf water potential, reduced chlorophyll, and other photosynthetic metabolites (Meyer et al. 2014; Lovell et al. 2016; Liu et al. 2015). Comparative analysis of rhizosheath metabolites further showed an increase in amino acids, carbohydrates and organic acids in response to drought (Liu et al. 2019).

In contrast, knowledge of the contribution of specialized metabolites to switchgrass stress response mechanisms has remained largely unexplored. For example, drought-induced alterations in terpenoid and phenylpropanoid metabolism have been reported (Meyer et al. 2014). In addition, our prior work revealed an expansive network of terpenoid-metabolic *terpene synthase* (*TPS*) and *cytochrome P450 monooxygenase* (*P450*) genes in *P. virgatum* var. Alamo, and combined metabolite and transcript profiling illustrated the formation of species-specific diterpenoids and the corresponding biosynthetic genes in switchgrass leaves and roots exposed to UV radiation and oxidative stress (Pelot et al. 2018; Muchlinski et al. 2019; Tiedge et al. 2020). Furthermore, emission of volatile mono- and sesqui-terpenoids was observed from switchgrass leaves and roots upon herbivore stress and treatment with defense-related plant hormones (Muchlinski et al. 2019). These collective insights support the importance of terpenoids and other specialized metabolite classes for switchgrass abiotic and biotic stress tolerance. Recent maize (*Zea mays*) and rice (*Oryza sativa*) studies showing induced diterpenoid formation under UV, oxidative and drought stress and decreased abiotic stress tolerance in diterpenoid-deficient maize mutants support a broader role of terpenoids in abiotic stress adaptation in monocot crops (Schmelz et al. 2014; Ding et al. 2021; Park et al. 2013; Horie et al. 2015; Vaughan et al. 2015a).

A deeper understanding of the relevance of specialized metabolism in drought tolerance and more broadly climatic adaptation in perennial biofuel crops can provide resources to improve breeding strategies for developing locally and broadly adapted feedstock systems (Morrow et al. 2014). In this study, we integrated tissue-specific transcriptome and metabolite analyses to investigate specialized metabolism responses to drought in two major switchgrass ecotypes with distinct habitats and contrasting drought tolerance (Liu et al. 2015), namely the lowland Alamo and the upland Cave-in-Rock genotypes.

## Results

To investigate metabolic drought responses in switchgrass, we selected the lowland genotype Alamo (AP13) and the upland genotype Cave-in-Rock which were ranked among the most drought-tolerant and drought-susceptible genotypes in a comparative study of 49 switchgrass varieties (Liu et al. 2015). At the beginning of the reproductive stage (R1), plants were exposed to four weeks of continuous drought treatment, whereby soil water content measured as Available Water Capacity (AWC) remained stable at 75% in well-watered control plants and decreased from 75% to 0% in drought-stressed plants (Supporting Information Fig. S1). Leaf and root tissue of both genotypes and treatment groups was harvested before the treatment (week 0), after two weeks and after four weeks, and samples were subject to transcriptomic and metabolomic analyses. Between three and four weeks of drought treatment Cave-in-Rock plants displayed an increasing wilting phenotype, whereas Alamo plants showed no or only minor wilting throughout the four-week drought treatment (Fig. 1).

**Fig. 1:**
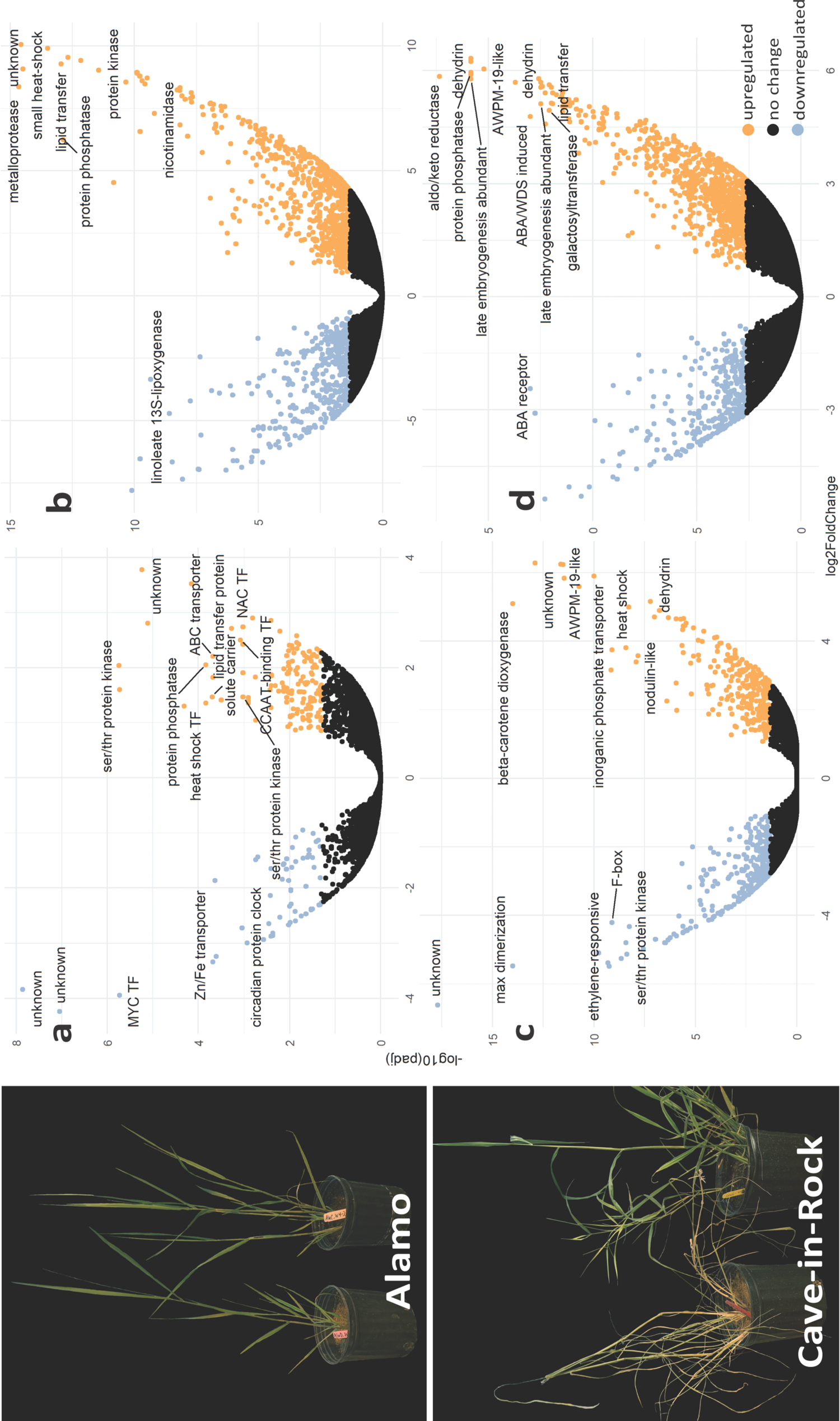
Left panel: Photographs of Alamo and Cave-in-Rock plants after four weeks of drought treatment (+d) or normal watering (-d). **Right panels:** Volcano plots of differentially expressed genes identified after four weeks of drought in (**a**) Alamo leaves (**b**) Cave-in-Rock leaves (**c**) Alamo roots (**d**) Cave-in-Rock roots. Differential expression threshold: padj < 0.05, log2 FC, and >1.

### Alamo and Cave-in-Rock plants show distinct transcriptomic alterations in response to drought

Illumina Novaseq 6000 RNA sequencing yielded a total of 2.4 billion and 2.7 billion high-quality reads for Alamo and Cave-in-Rock samples, respectively, representing >97% of the total reads obtained in both datasets. Alignment of the high-quality sequences against the switchgrass Alamo AP13 genome (phytozome-next.jgi.doe.gov/info/Pvirgatum_v5_1) resulted in average mapping rates of 86% for Alamo and 83% for Cave-in-Rock, thus providing a comprehensive transcriptomic dataset for gene discovery and gene expression analyses. Using this dataset, differential gene expression analysis was performed for control and drought-treated plants of both genotypes. A total of 565 differentially expressed genes (DEGs) were identified in roots and 204 DEGs in leaves (DEG threshold: padj < 0.05; |log2 FC| > 1) of Alamo plants after four weeks of drought treatment compared to well-watered control plants (Fig. S2). Cave-in-Rock plants showed stronger drought-induced changes with a total of 1198 DEGs in roots and 1120 DEGs in leaves; constituting 2- and 5-fold more DEGs as compared to Alamo roots and leaves, respectively (Fig. S2). In addition, Cave-in-Rock plants showed an earlier onset of transcriptomic changes compared to Alamo. After two weeks of watering withdrawal, 2951 and 897 genes were differentially expressed in Cave-in-Rock leaves and roots, respectively, whereas only 19 and 128 genes were differentially expressed in Alamo leaves and roots (Supporting Information Table S1), concurrent with the earlier onset of phenotypic drought symptoms in Cave-in-Rock (Fig. S2). Furthermore, differential gene expression was more pronounced in roots than leaves in both genotypes. Indeed, permutational multivariate analysis of variance (PERMANOVA) illustrated that tissue type (leaves or roots) had the predominant impact on gene expression levels (38.6%, *p*<0.001***), followed by genotype (8%, *p*<0.003**) and stress treatment (6%, *p*<0.007**) (Supporting Information Table S2).

### Major transcriptomic changes include genes of known drought response mechanisms and core general metabolic processes

Consistent with the differences in the number and onset of DEGs between Alamo and Cave-in-Rock, the identified genes showing the most significant differential expression differed between the two genotypes and tissues (Fig. 1). Only two genes, a putative circadian clock protein (*Pavir.9NG553200*) and a predicted lipid transfer protein (*Pavir.5NG603383*) were differentially expressed in all genotypes and tissues (Fig. S2). The genotype- or tissue-specific DEGs included numerous so far uncharacterized genes as well as several genes associated with known drought response mechanisms. For example, in roots of both genotypes *dehydrin* (log_2_(fold change) Alamo=5.2, CiR=6.24) and other *Late Embryogenesis Abundant* (*LEA*) genes (log_2_(fold change) Alamo=4.34, CiR=5.78), as well as several *AWPM19-like* plasma-membrane-associated abscisic acid (ABA) influx transporters implicated with drought tolerance (log_2_(fold change) Alamo=6.25, CiR=6.04) were highly upregulated (Fig. 1, Supporting Information Table S1). Notably, among the 11 *AWPM19-like* genes, Alamo and Cave-in-Rock featured distinct genes (*Pavir.2NG274300* in Alamo versus *Pavir.9NG018700* in Cave-in-Rock) as the most differentially expressed *AWPM19-like* genes. The increased expression of known drought-associated genes supports the drought response in both genotypes, despite the visual lack of a wilting phenotype in Alamo plants.

Next, we investigated the impact of drought on core metabolic pathways independent of known drought response processes. Among 26 GO terms significantly enriched in all samples combined most encoded for biological processes or molecular functions (Fig. S3). In Alamo leaves DNA-binding and electron carrier processes were most significantly enriched, whereas carbohydrate metabolism was predominant in roots. Interestingly, Cave-in-Rock roots featured highly enriched abiotic stress response processes rather than carbohydrate metabolism under drought stress, whereas leaves showed patterns similar to Alamo with DNA-binding and hydrolase activities being differentially expressed (Fig. S3). Additional pathway enrichment analyses using KEGG terms confirmed a substantially higher number of metabolic pathways enriched in Cave-in-Rock as compared to Alamo, with more genes underlying the enriched pathways on average (Fig. 2 and Fig. S4). Plant signal transduction process ranked among the most significantly enriched in Alamo and Cave-in-Rock leaves, especially in response to drought stress. By contrast, endoplasmic reticulum protein processing and nitrogen metabolism were most significantly altered in Alamo and, to lesser degree, in Cave-in-Rock roots (Fig. 2). Albeit at lower levels, pathway enrichment was also observed for general and specialized metabolism, including carbon and amino acid metabolism, as well as carotenoid, steroid, and phenylpropanoid biosynthesis (Fig. 2, Supporting Information Table S3).

**Fig. 2:**
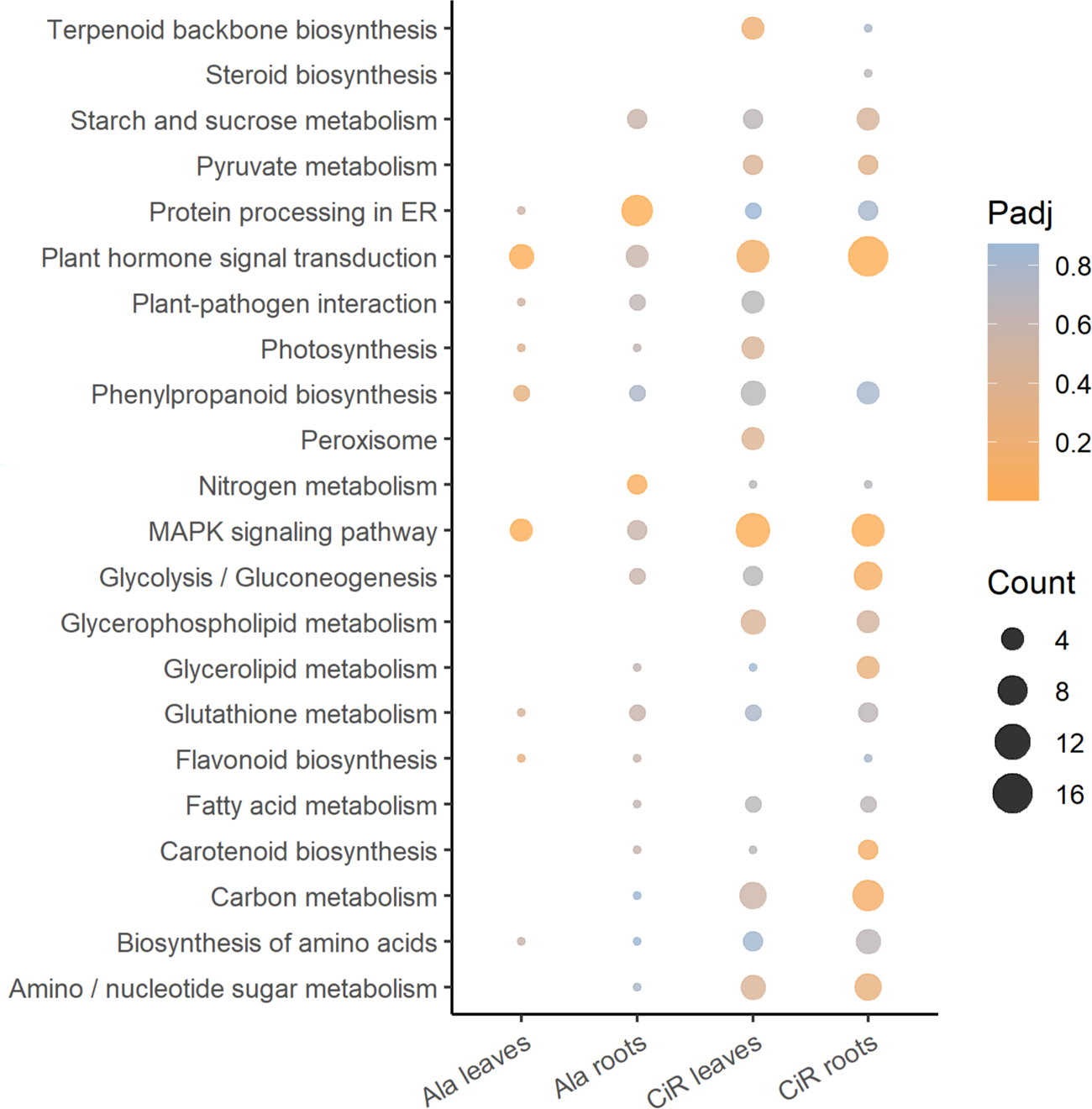
KEGG pathway enrichment analysis of differentially expressed genes. Circle color denotes the adjusted *p*-value (Padj), circle size is proportional to the number of genes involved in the enrichment of the pathway (Count).

### Switchgrass features tissue-, genotype- and drought-specific alterations in terpenoid and phenylpropanoid pathways

The enrichment of terpenoid and phenylpropanoid metabolism is consistent with prior studies illustrating the up-regulation, albeit at moderate levels, of switchgrass terpenoid- and phenylpropanoid-metabolic pathways in response to drought, UV irradiation and oxidative stress (Meyer et al. 2014; Pelot et al. 2018; Muchlinski et al. 2019; Tiedge et al. 2020). To investigate in more detail the impact of drought on switchgrass specialized metabolism, we compared the transcript abundance of key genes of the terpenoid and phenylpropanoid scaffold-forming pathways in both genotypes. Due to the lack of a Cave-in-Rock genome, gene annotations are based on homology searches against the switchgrass Alamo AP13 genome (phytozome-next.jgi.doe.gov/info/Pvirgatum_v5_1). Interestingly, the focal genes showed similar tissue-specific expression profiles in Alamo and Cave-in-Rock and no substantial drought-induced gene expression changes were observed (Fig. 3). For example, of the four annotated *1-deoxyxylulose 5-phosphate synthase* (*DXS*) genes of the methylerythritol phosphate (MEP) pathway, two homologs, *Pavir.3KG128241* and *Pavir.3NG140939*, were abundant in leaves but ∼10-60-fold less in roots of both Alamo and Cave-in-Rock (Fig.3a). Likewise, the predicted squalene synthase, *Pavir.4NG340500*, displayed high abundance in leaves and low gene expression in roots. However, select genes showed genotype-specific differences in their expression patterns. This included the predicted *geranylgeranyl pyrophosphate synthase* (*GGPPS*), *Pavir.6Ng089000*, that was expressed in Alamo but not Cave-in-Rock roots (Fig. 3a). A similar trend of gene expression was observed for core genes of phenylpropanoid metabolism with several annotated *phenylalanine ammonia lyase* (*PAL*), *cinnamate-4-hydroxylase* (*C4H*), and *4-coumaroyl-CoA ligase* (*4CL*) genes showing higher expression in roots as compared to leaves in both Alamo and Cave-in-Rock (Fig. 3b).

**Fig. 3:**
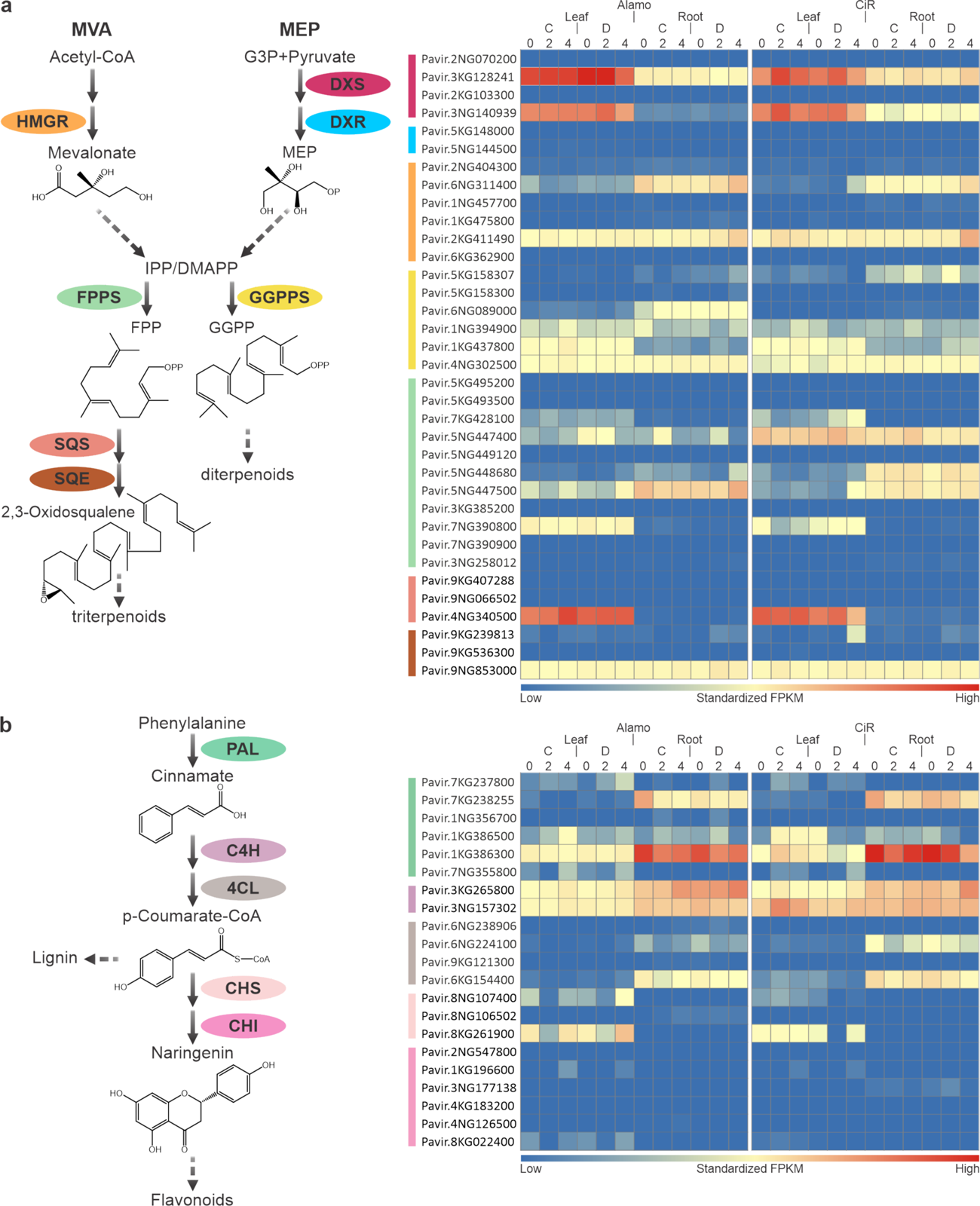
Plot of normalized gene expression profiles of genes with predicted functions in (**a**) terpenoid backbone biosynthesis and (**b**) flavonoid backbone biosynthesis after 0, 2 or 4 weeks in drought-treated (D) or well-watered control (C) Alamo and Cave-in-Rock (CiR) plants. Gene expression data are based on four biological replicates and gene functional annotations are based on best matches in BLAST searches against public and in-house protein databases. Gene IDs were derived from the *Panicum virgatum* genome v5.1 (phytozome-next.jgi.doe.gov/info/Pvirgatum_v5_1). Abbreviations: *DXS, 1-deoxyxylulose 5-phosphate synthase; DXR, 1-deoxyxylulose 5-phosphate reductase; HMGR, HMG-CoA reductase; FPPS, farnesyl pyrophosphate synthase; GGPPS, geranylgeranyl pyrophosphate synthase; SQS, squalene synthase; SQE, squalene epoxidase; PAL, phenylalanine ammonia lyase; C4H, cinnamate-4-hydroxylase; 4CL, 4-coumaroyl-CoA ligase; CHS, chalcone synthase; CHI, chalcone isomerase*.

Contrasting the largely unaltered and comparable expression of the highly conserved upstream pathway genes, both Alamo and Cave-in-Rock plants featured distinct gene expression profiles for downstream pathway branches that generate species-specific, functionalized metabolites. Following the recent discovery of specialized triterpenoid and steroid saponins in switchgrass (Li et al. 2020), identification and gene expression analysis of predicted *cycloartenol synthases* (*CAS*), *lanosterol synthases* (*LAS*), *β-amyrin synthases* (*BAS*), as well as members of the *CYP71A*, *CYP90B* and *CYP94D* cytochrome P450 families and *sterol 3-β-glucosyltransferases* with known functions in triterpenoid metabolism revealed distinct expression patterns across tissue type and genotype (Fig. 4). For example, hierarchical gene cluster analysis illustrated predicted *triterpenoid synthase* (*TTS*) genes and a putative *CYP72A* gene with similar inducible expression patterns in Alamo leaves after two and four weeks of drought (Supporting Information Table S4). Likewise, in Alamo roots a different group of *TTS*, *sterol 3-β-glucosyltransferase*, and putative *CYP72A*, *CYP94D* and *CYP90B* genes, known to function in the biosynthesis of triterpenoid saponins such as diosgenin (Ciura et al. 2017), displayed common inducible expression patterns after four weeks of drought (Fig. 4 upper panel). By contrast, co-expression patterns of triterpenoid-biosynthetic genes were not detectable in Cave-in-Rock (Fig. 4 lower panel).

**Fig. 4:**
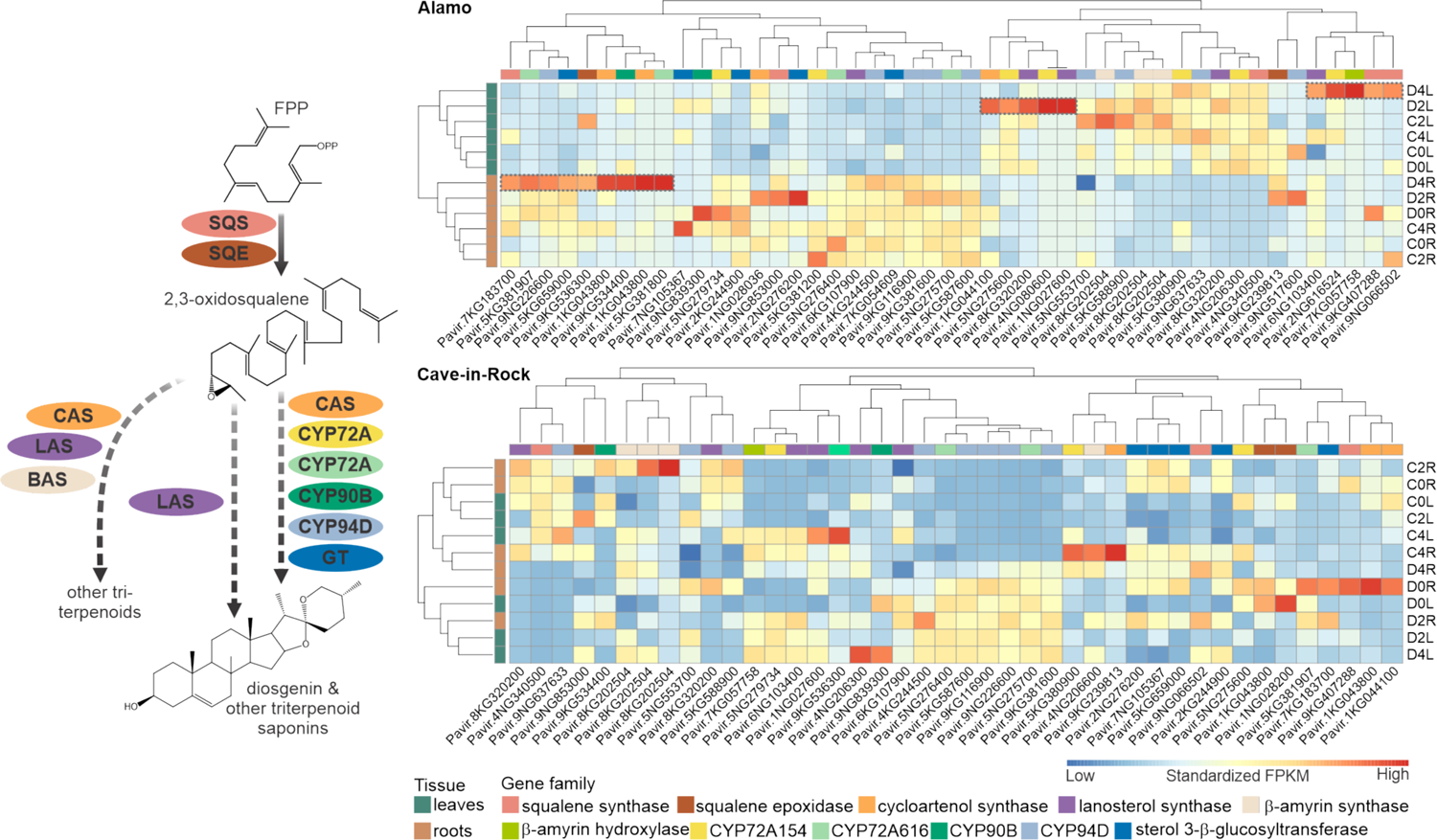
Hierarchical cluster analysis of select genes with predicted functions in triterpenoid biosynthesis in Alamo and Cave-in-Rock. Gene functional annotations are based on best matches in BLAST searches against in-house protein databases of known triterpenoid-metabolic genes. Gene IDs are derived from the *Panicum virgatum* genome v5.1 (phytozome-next.jgi.doe.gov/info/Pvirgatum_v5_1). Gene expression data are based on four biological replicates. Dashed boxes highlight genes with relevant co-expression patterns. Right side: C0L, C2L, C4L: Leaves of well-watered control plants after 0, 2 and 4 weeks of treatment; C0R, C2R, C4R: Roots of well-watered control plants; D0L, D2L, D4L: Leaves of drought-stressed plants; D0R, D2R, D4R: Roots of drought-stressed plants.

Our prior research identified expansive, species-specific diterpene synthase (diTPS) and P450 families in switchgrass that form complex metabolic networks toward a range of labdane-related diterpenoids, including *syn*-pimarane and furanoditerpenoid compounds that occur, perhaps uniquely, in switchgrass (Fig. S5) (Pelot et al. 2018; Muchlinski et al. 2021). This pathway knowledge enabled a detailed analysis of transcriptomic alterations related to diterpenoid metabolism. Contrasting the largely similar expression patterns of MEP and MVA pathway genes (Fig. 3a), hierarchical gene cluster analysis revealed distinct *diTPS* and *P450* gene expression between Alamo and Cave-in-Rock (Fig. 5). In Alamo roots, the *cis-trans-clerodienyl pyrophosphate (CLPP) synthase PvCPS1* and the *P450* genes, *CYP71Z25*, *CYP71Z26*, and *CYP71Z28* shown to form furanoditerpenoids (Muchlinski et al. 2021), showed patterns of co-expression at two and four weeks of drought. Similarly, the predicted *syn-CPP synthases*, *PvCPS9* and *PvCPS10*, as well as two class I diTPS, *PvKSL4* and *PvKSL5*, shown to form *syn*-pimaradiene compounds (Pelot et al. 2018), co-expressed in roots, albeit without significant drought-inducible transcript changes. In Cave-in-Rock, *diTPS* and *P450* genes were expressed mostly in the well-watered plants and water deficiency did not elicit significant transcript accumulation. Also, contrasting roots, no drought-elicited changes in the expression of diterpenoid pathway genes was detectable in leaves of either ecotype.

**Fig. 5:**
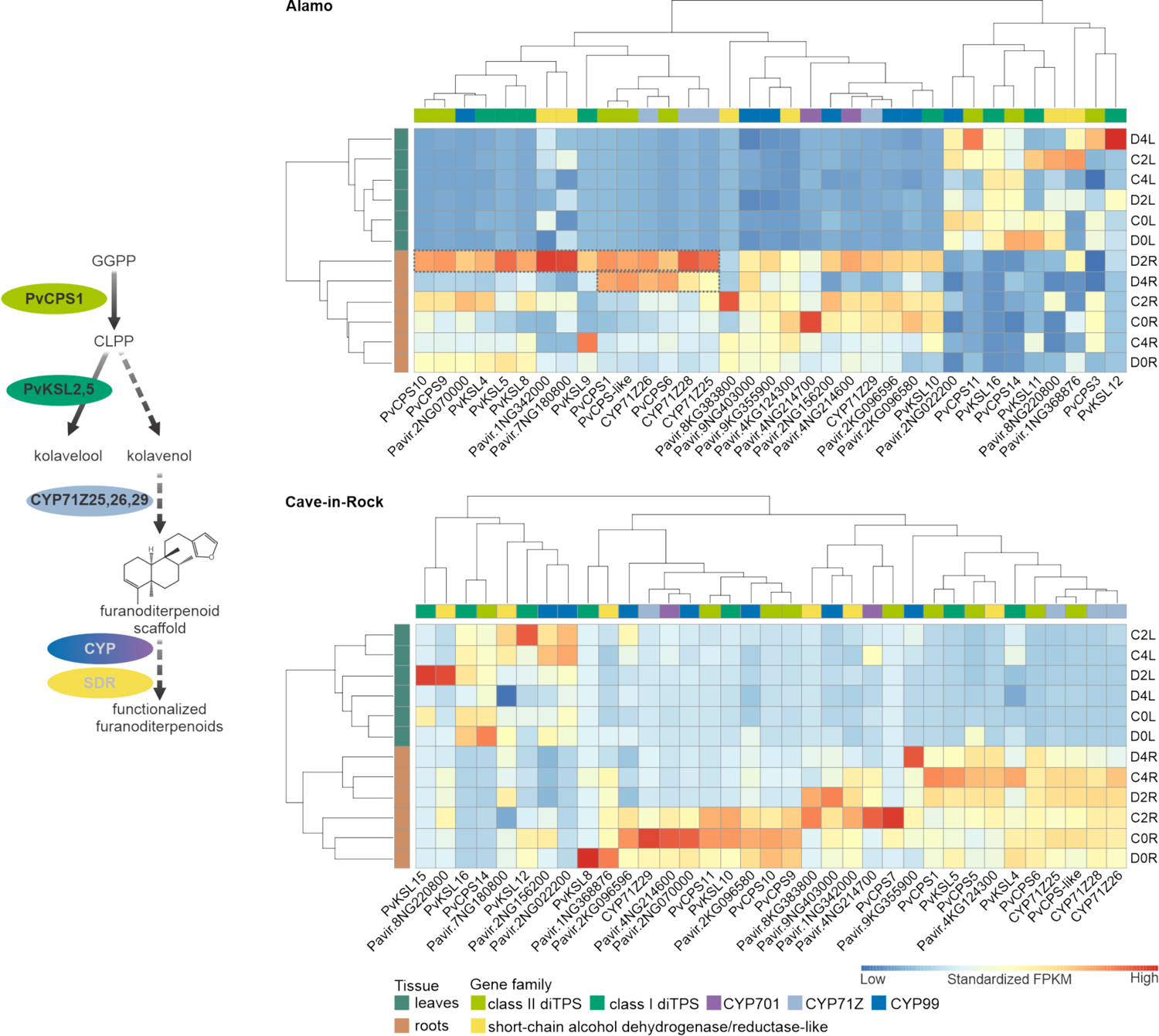
Hierarchical cluster analysis of select genes with known or predicted functions in diterpenoid biosynthesis in Alamo and Cave-in-Rock. Gene functional annotations are based on previous biochemical enzyme characterizations or best matches in BLAST searches against in-house protein databases of known diterpenoid-metabolic genes. Gene IDs are derived from the *Panicum virgatum* genome v5.1 (phytozome-next.jgi.doe.gov/info/Pvirgatum_v5_1). Gene expression data are based on four biological replicates. Dashed boxes highlight genes with relevant co-expression patterns. C0L, C2L, C4L: Leaves of well-watered control plants after 0, 2 and 4 weeks of treatment; C0R, C2R, C4R: Roots of well-watered control plants; D0L, D2L, D4L: Leaves of drought-stressed plants; D0R, D2R, D4R: Roots of drought-stressed plants.

### Switchgrass leaves and roots show drought-inducible metabolite alterations

To complement transcriptomic studies leaf and root metabolomes of Alamo and Cave-in-Rock plants under drought-stressed and well-watered conditions were examined using untargeted liquid-chromatography-quadrupole Time of Flight-mass spectrometry (LC-QToF-MS) analysis. Using accurate mass, retention time (RT), and fragmentation patterns, we identified 5181 and 3234 metabolite features in positive and negative ion mode, respectively. To compare metabolite profiles across tissues, genotypes and treatments, after filtering for ion-abundance (see Methods) 2519 positive mode mass features (identified as retention time-*m*/*z* ratio pairs) were selected for downstream statistical analysis (Supporting Information Table S5).

Aligned with the transcriptomic changes, biostatistical analysis of the untargeted metabolomic data via PERMANOVA showed that tissue type had the highest impact on metabolite composition (72.4%, *p*<0.001***), followed by difference in genotype (2.8%, *p*<0.028*) and drought-treatment versus control (0.6%, *p*<0.332) (Supporting Information Table S2). Hence, we further analyzed the metabolite profiles independently within each tissue type. Despite the relatively lower impact of drought treatment on metabolic alterations, under a multivariate dimension-reduction based on genotype metabolite features clustered together prior to water deprivation (week 0), but separated in leaves and, to a lesser extent, in roots after four weeks of drought treatment (Fig. 6). This shift in metabolite composition was driven by several major features (Fig. 7). In leaves, most compounds showing accumulation differences in control and drought-stressed plants were annotated as phospholipids based on database searches for each feature. These compounds increased in abundance in Alamo during drought treatment, whereas a decrease was observed in Cave-in-Rock (Fig. 7). In addition to predicted phospholipids, a few features were enriched in Alamo leaves under drought stress, whereas many unidentified compounds were enriched in drought-treated Cave-in-Rock leaves (Supporting Information Table S2). In contrast, several compounds identified as diterpenoids and triterpenoids by comparison of RT and fragmentation patterns to previously identified compounds (Li et al., 2020; Muchlinski et al., 2021) accumulated in roots of both Alamo and Cave-in-Rock plants under drought stress, with a stronger increase in Alamo (Fig. 7). Other root metabolites that accumulated differentially under drought conditions either did not score significant database matches or could only be assigned to the general classes of carbohydrates, acids, or alcohols (Fig. 7). Among the few features that were generally enriched in both leaves and roots and in both ecotypes under water deficient conditions was also abscisic acid (ABA), which was increased by ∼40-300-fold under drought (Fig. 8a, Supporting Information Table S6, ABA: 6.08_247.1244m/z).

**Fig. 6:**
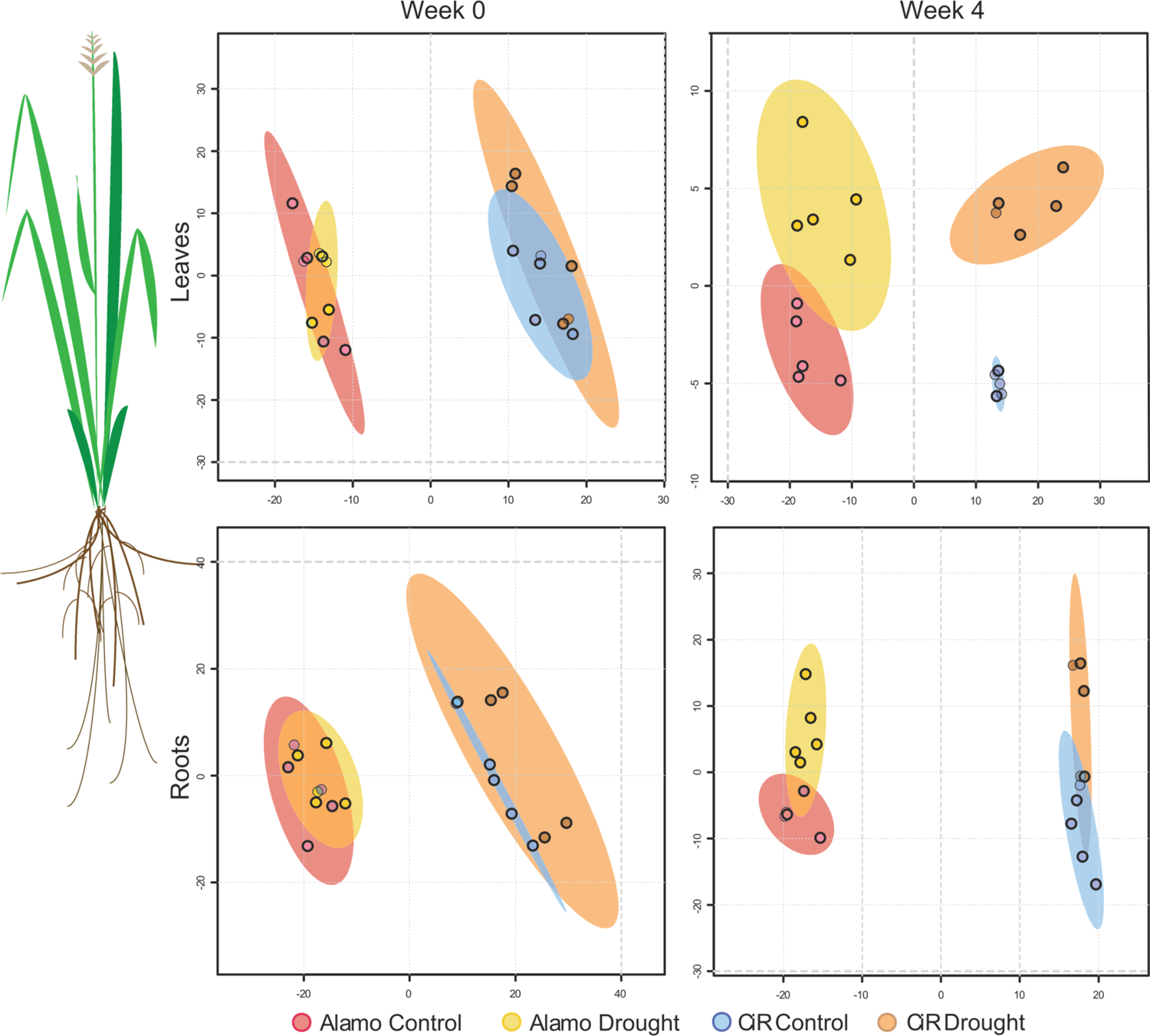
Partial Least-Squares Discriminant Analysis (PLS-DA) plots of LC-MS positive mode metabolome divergence based on five biological replicates. X-axis: Principal Component 1; y-axis: Principal Component 2.

**Fig. 7:**
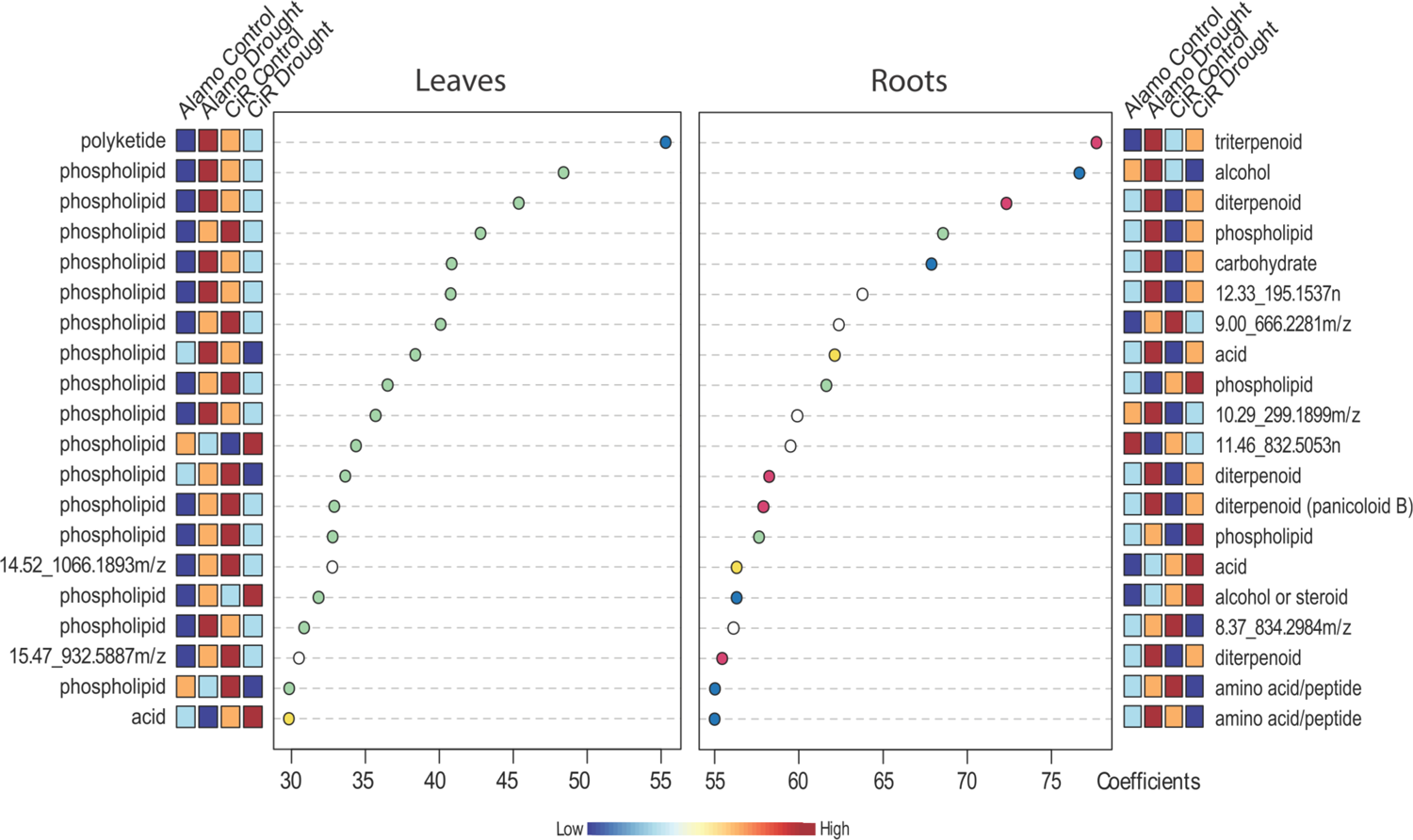
Scree plot of metabolite features obtained via LC-MS positive mode analysis that show the most significant contribution to changes in metabolite profiles in response to drought stress. A higher coefficient (x-axis) denotes a higher importance for this feature in the Partial Least-Squares Discriminant Analysis (PLS-DA) shown in Fig. 6. Boxes display the relative abundance of a feature among the different groups as based on five biological replicates. Metabolite annotations are based on matching *m*/*z* ratios, RT and fragmentation patterns against online databases.

**Fig. 8:**
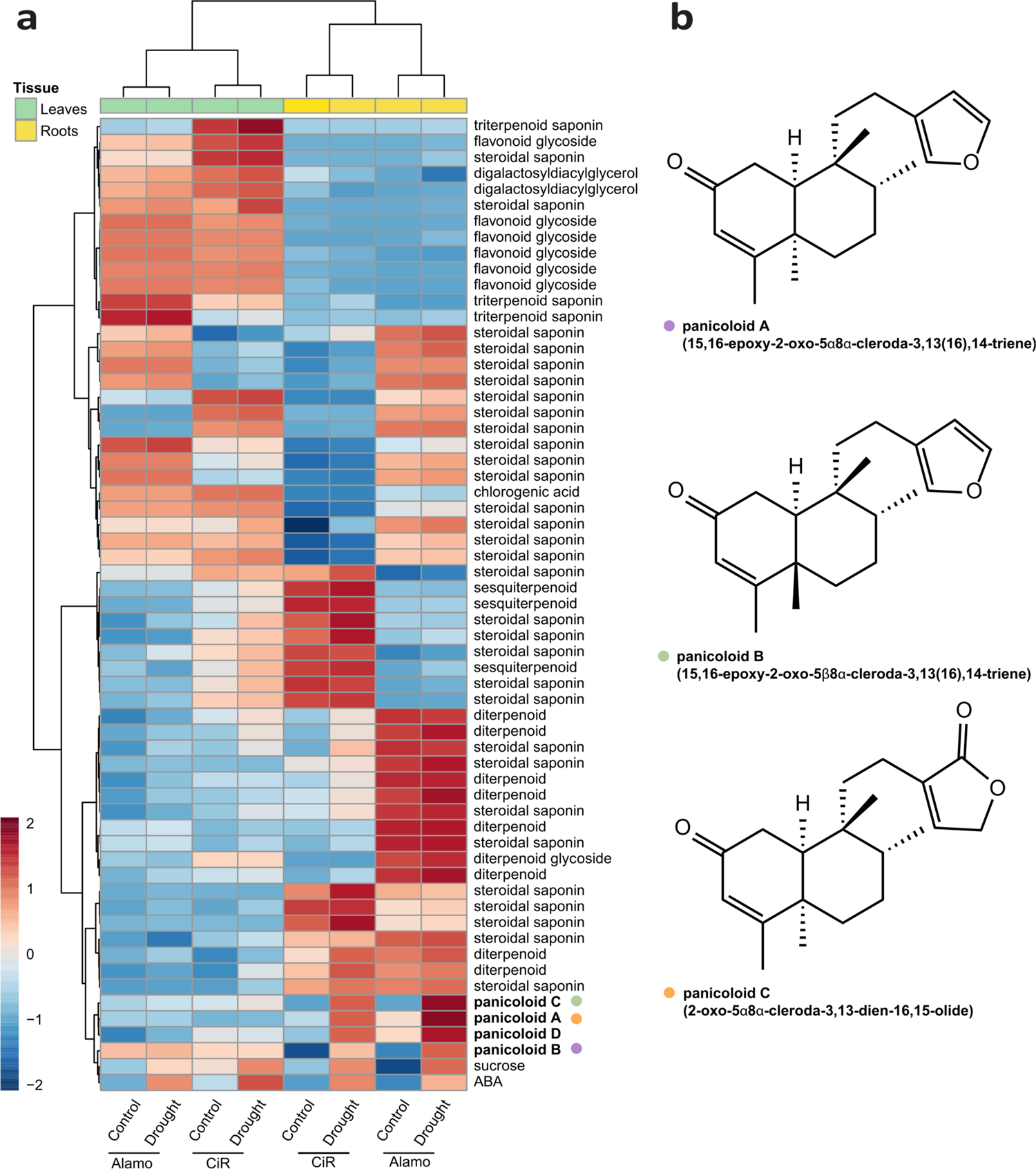
(**a**) Hierarchical cluster analysis of select specialized metabolite accumulation patterns. Sucrose and abscisic acid (ABA) abundance provide as drought-related reference metabolites. Metabolite annotations are based on matching *m*/*z* ratios, RT and fragmentation patterns against online databases. (**b**) Structures of drought-induced diterpenoids, 15,16-epoxy-2-oxo-5*α*8*α*-cleroda-3,13(16),14-triene (*m*/*z* 301, panicoloid A), 15,16-epoxy-2-oxo-5β8*α*-cleroda-3,13(16),14-triene (*m*/*z* 301, panicoloid B) and 2-oxo-5*α*8*α*-cleroda-3,13-dien-16,15-olide (*m*/*z* 317, panicoloid C), isolated from drought-stressed switchgrass roots. Metabolite abundance is based on five biological replicates.

### Drought induces the accumulation of specialized furanoditerpenoids in switchgrass roots

Considering that predicted specialized steroidal and triterpenoid saponins and diterpenoids contribute substantially to the metabolic differences between Alamo and Cave-in-Rock roots, we examined these compounds in more detail in leaf and root extracts after four weeks of drought where the physiological stress symptoms were most pronounced. Several predicted flavonoid glycosides were identified in leaves of Alamo and Cave-in-Rock plants but were absent in root tissue. However, these compounds showed no or only minimal patterns of drought-elicited accumulation (Fig. 8a). In addition, a range of distinct terpenoid metabolites was identified. Among these metabolites, the largest group represented recently identified steroidal or triterpenoid saponins (Li et al. 2020), showing distinct profiles across tissues and genotypes. Triterpenoid saponins occurred predominantly in leaves of both Alamo and Cave-in-Rock plants, whereas the larger group of steroidal saponins were present in leaves and/or roots and occurred predominantly in either Alamo or Cave-in-Rock plants. Despite the overall abundance of these saponins, the vast majority of the annotated metabolites were not significantly enriched upon drought stress (p ≤ 0.05, Supporting Information Table S6). In addition to the larger group of triterpenoids, 11 compounds predicted as specialized diterpenoids were identified, the majority of which occurred predominantly in Alamo roots and were absent or abundant at only low levels in Cave-in-Rock (Fig. 8a). Notably, two pairs of predictably isomeric diterpenoids were detected in Alamo and Cave-in-Rock that showed substantial accumulation mostly in drought-stressed roots. One metabolite pair at retention times of 9.06 min and 9.26 min featured a dominant precursor mass ion of *m*/*z* 317 [M+H]^+^ and one compound pair at retention times of 11.50 min and 11.72 min featured a dominant ion of *m*/*z* 317 [M+H]^+^. Together with the presence of additional mass ions of *m*/*z* 257, *m*/*z* 189, *m*/*z* 177 or *m*/*z* 135, these fragmentation patterns suggested that these compounds represent labdane-related diterpenoids carrying one or more oxygenation functions (Supporting Information Fig. S4). To elucidate the precise structure of these diterpenoids, metabolites were extracted from mature Cave-in-Rock roots and purified via liquid-liquid phase partitioning followed by HPLC. Purified samples (0.4-0.8 mg for each compound) of both *m/z* 301 isomers and both *m/z* 317 diterpenoid isomers were used for 1D (^1^H and ^13^C) and 2D (HSQC, COSY, HMBC and NOESY) NMR analyses. Collectively, the generated data identified the *m*/*z* 301 diterpenoids as 15,16-epoxy-2-oxo-5*α*8*α*-cleroda-3,13(16),l4-triene and its C19 enantiomer, while the earlier eluting diterpenoid *m*/*z* 317 was identified as 2-oxo-5*α*8*α*-cleroda-3,13-dien-16,l5-olide, together designated as panicoloid A,B, and C respectively (Fig. 8b, Supporting Information Fig. S6). Insufficient abundance and purity of the later eluting *m*/*z* 317 diterpenoid isomer prevented structural analysis of this metabolite. Based on the similar mass fragmentation pattern and retention time compared to the other *m*/*z* 317 isomer (Supporting Information Fig. S4), it is plausible that this compound represents its enantiomeric isomer and is tentatively named panicoloid D. The identified diterpenoids represent derivatives of previously identified switchgrass furanoditerpenoids that feature additional carbonyl functions at C2 and/or C14 and are collectively named here as the group of panicoloids (Fig. S4).

## Discussion

Knowledge of the gene-to-metabolite relationships underlying specialized metabolic pathways that contribute to plant stress resilience can enable new crop optimization strategies for addressing exacerbating environmental pressures and associated harvest loss (Savary et al. 2019). Extensive studies in major food and bioenergy crops such as rice, maize, and sorghum (*Sorghum bicolor*) have demonstrated that species-specific blends of specialized terpenoids, phenylpropanoids, oxylipins, and benzoxazinoids mediate complex responses to biotic and abiotic perturbations (Schmelz et al. 2014; Murphy and Zerbe 2020). By contrast, knowledge of the diversity of specialized metabolites in the perennial bioenergy crop switchgrass and its relevance for drought adaptation in different switchgrass genotypes is incomplete. Combined transcriptome and metabolome analysis of the lowland ecotype Alamo shown to be drought-tolerant and the upland ecotype Cave-in-Rock with low drought tolerance (Liu et al. 2015) revealed common and distinct metabolic alterations and identified specialized diterpenoid metabolites with possible functions in switchgrass drought adaptation.

Consistent with prior studies showing that upland and lowland switchgrass ecotypes have different transcriptomic profiles under optimal conditions (Ayyappan et al. 2017), this study demonstrates that metabolic alterations in Alamo and Cave-in-Rock are predominantly driven by differences in tissue type and genotype, thus reflecting the different habitat range and climatic adaptation of switchgrass ecotypes. A stronger wilting phenotype after four weeks of drought treatment, along with more than twice as many differentially expressed genes illustrate more pronounced drought-induced metabolic changes in Cave-in-Rock, consistent with prior studies identifying Cave-in-Rock as particularly drought-susceptible among switchgrass varieties (Liu et al. 2015). Presence of known drought-response genes among the most differentially expressed genes in Alamo and/or Cave-in-Rock supported substantial drought responses in both genotypes during drought treatment, despite the lack of significant phenotypic changes in Alamo. Know drought-associated genes included *LEA* genes, including dehydrin shown to impact drought tolerance in Arabidopsis and cotton (*Gossypium spec.*) (Olvera-Carrillo et al. 2010; Magwanga et al. 2018), a NAC transcription factor (*Pavir.8KG003520*) shown to contribute to drought tolerance in a recent switchgrass GWAS study (Lovell et al. 2021), and AWPM-19-like abscisic acid (ABA) influx transporters (Yao et al. 2018), Differential expression of several AWPM-19-like genes in both Alamo and Cave-in-Rock is consistent with an increase in ABA observed in both genotypes and tissues in response to drought stress. While prior studies that showed increased ABA and sugar accumulation in drought-tolerant switchgrass genotypes (Liu et al. 2015), the final concentration of ABA in drought stressed plants was at comparable levels in both genotypes in our study, whereas the concentration in the control plants was lower in Alamo, resulting in a higher fold-change/increase in Alamo when compared to Cave-in-Rock Notably, the specific genes of the above gene families being differentially expressed differed between Alamo and Cave-in-Rock, suggesting that switchgrass genotypes recruit specific genes governing stress response mechanisms.

Pathways of general and specialized metabolism showed overall comparatively moderate differential gene expression in response to drought stress with apparent metabolic differences between leaf and root tissue. Accumulation of sucrose and ABA in Alamo and Cave-in-Rock tissues is consistent with previously demonstrated switchgrass drought responses (Liu et al. 2015). In addition, the observed major contribution of phospholipids to metabolic alterations in leaves of both genotypes may be related to an upregulation of pathways involved in membrane lipid systems and cell wall biosynthesis as shown in, for example, drought-resistant maize lines (Zhang et al. 2020). Additional phospholipid roles in drought tolerance may include maintenance of membrane integrity or mitigation of drought-related cell damage (Hamrouni et al. 2001), as well as signaling processes during water deficiency as reported in selected drought-resistant species (Moradi et al. 2017; Quartacci et al. 1995). Contrasting drought-related flavonoid functions in, for example, wheat (*Triticum aestivum*) and other species (Gai et al. 2020; Ma et al. 2014), leaf flavonoid glycosides did not accumulate in response to drought in either switchgrass genotype, despite a moderate upregulation of select pathway genes such as *CHS* in drought-stressed plants. Similarly, steroidal and triterpenoid saponins were identified in switchgrass leaves, and have been shown to increase as part of the leaf cuticular waxes in response to drought (Kim et al. 2007). However, our metabolite analysis did not show significant drought-elicited triterpenoid accumulation that would support a similar function in switchgrass.

Different from leaves, steroidal and other triterpenoid saponins as well as diterpenoids constituted the major determinants of metabolic differences in drought-stressed roots. Notably, MEP and MVA pathway genes showed no or very minor patterns of drought-inducible expression in Alamo or Cave-in-Rock, indicating that no major change in the production of terpenoid precursors occurs in response to drought. One exception is the putative *GGPPS*, *Pavir.6NG089000*, that is expressed in Alamo but not Cave-in-Rock roots and may contribute to differences in terpenoid metabolism between these genotypes. This apparent lack of drought activation of terpenoid backbone pathways suggests that terpenoid accumulation has to derive from existing precursor pools via changes in pathway branches en route to specific terpenoids. Indeed, in leaves and roots of Alamo, but not Cave-in-Rock, downstream triterpenoid-metabolic pathway genes increased under drought conditions. Interestingly, no apparent triterpenoid-biosynthetic gene clusters were identified in the switchgrass genome, which differentiates switchgrass from other plant species where triterpenoid metabolism is commonly arranged in form of often stress-inducible biosynthetic clusters (Liu et al. 2020; Bai et al. 2021). Despite the lack of apparent genomic clusters, clear co-expression patterns were observed for several *TTS* genes as well as *sterol 3-β-glucosyltransferases* and *P450* genes of the *CYP71A*, *CYP94D* and *CYP90B* families, thus supporting the presence of co-expressed pathways toward specific triterpenoids. The identification of diverse mixtures of diosgenin and closely related triterpenoid saponins in roots of several upland and lowland ecotypes supports this hypothesis (Lee et al. 2009; Li et al. 2020). However, the annotated triterpenoid metabolites were not significantly enriched in response to drought in either genotype, suggesting that the inducible expression of triterpenoid pathways in Alamo is related to distinct drought-stress responses. It can be speculated that root triterpenoids serve antioxidant functions to mitigate oxidative damage caused by water deficiency as shown in Arabidopsis and other species (Posé et al. 2009; Nasrollahi et al. 2014; Puente-Garza et al. 2017). Also, recent studies have demonstrated bioactive triterpenoids in roots exudates of soybean (*Glycine max*) and tomato (*Solanum lycopersicum*) that aid the assembly of the root microbiome to confer robustness against environmental stresses (Fujimatsu et al. 2020; Nakayasu et al. 2021). While distinct microbiome responses to drought stress have been reported in upland and lowland switchgrass ecotypes (Liu et al. 2021), the role of specialized metabolites such as saponins in these interactions is yet to be discovered.

Unlike root triterpenoids, significant drought-induced accumulation of several diterpenoids in roots supports a role in drought response mechanisms. Although requiring further biological studies, the higher abundance of these diterpenoids in Alamo as compared to Cave-in-Rock may contribute to the distinct stress resilience in these genotypes. In addition, accumulation in both Alamo and Cave-in-Rock support a role of these metabolites in drought responses rather than general stress responses associated with the more pronounced wilting phenotype observed in Cave-in-Rock. Structural analysis identified three compounds as oxygenated clerodane furanoditerpenoids, named here panicoloid A-C, which likely represent derivatives of furanoditerpenoid scaffolds recently identified in switchgrass (Pelot et al. 2018; Muchlinski et al. 2021). Notably, the enantiomeric stereochemistry of panicoloid B and C is likely derived from the activity of yet unidentified diTPS functionally related to the CLPP synthase PvCPS1. Drought-induced gene expression increases of characterized pathway genes toward clerodane-type furanoditerpenoids, including the diTPS *PvCPS1* and the P450 genes *CYP71Z25*, *CYP71Z26* and *CYP71Z28* supports a role of these pathway genes in panicoloid biosynthesis. Additional co-expression of select P450s of the CYP99 family and predicted short-chain alcohol dehydrogenases/reductases, shown to function in specialized diterpenoid metabolism in maize and rice (Swaminathan et al. 2009), suggests possible functions in the position-specific oxygenation reactions toward panicoloid biosynthesis. Similar to triterpenoid metabolism, the lack of co-expression of these genes in the drought-susceptible Cave-in-Rock genotype may support a role of panicoloids in switchgrass drought responses. Combined with the abundance of yet unidentified diterpenoids in switchgrass roots, drought-elicited co-expression patterns of additional predicted *syn-CPP synthase* genes (*PvCPS9, PvCPS10*) and characterized class I diTPS forming *syn*-pimarane diterpenoids (*PvKSL4, PvKSL5*) suggest the presence of a broader diversity of drought-induced diterpenoids in switchgrass. Similar clerodane-type furanoditerpenoids have also been identified in species of *Vellozia spec.* (Pinto et al. 1994), *Solidago spec.* (Anthonsen et al. 1973; McCrindle et al. 1976), and *Croton campestris* (El Babili et al. 1998), where they will have evolved independently given the phylogenetic distance between these plant genera. While the drought-induced expression of diterpenoid-metabolic genes and associated accumulation of panicoloids and possibly other diterpenoids supports a role in switchgrass drought tolerance, the underlying mechanisms will require future investigation. However, drought-related diterpenoid bioactivities have recently been supported in other monocots. For example, maize studies demonstrated the accumulation of specialized kauralexin and dolabralexin diterpenoids in response to oxidative, drought and salinity stress (Christensen et al. 2018; Mafu et al. 2018), and diterpenoid-deficient maize mutants show decreased resilience to abiotic stress (Vaughan et al. 2015b). Additionally, antioxidative functions in relation to drought stress have been shown for select diterpenoids (Munné-Bosch and Alegre 2003), and diterpenoid roles in the root microbiome assembly have been suggested based on changes in the microbiome composition in the kauralexin- and dolabralexin-deficient maize *an2* mutant (Murphy et al. 2021).

Collectively, these findings exemplify the power of combining transcriptomic, metabolomic and metabolite structural approaches to accelerate the discovery of plant specialized pathways and products to better understand their relevance and role in plant stress responses. This approach revealed common and distinct drought-induced metabolic changes in switchgrass genotypes of contrasting drought tolerance. These insights provide the foundation for future targeted genetic studies to investigate the diversity and protective function of terpenoids and other specialized metabolites in switchgrass drought tolerance.

## Methods

### Plant material and treatment

Switchgrass genotypes Alamo AP13 and Cave-in-Rock were kindly provided by Dr. Malay Saha (Noble Research Institute, USA). Plants were propagated from tillers to maintain low genetic variation and cultivated in greenhouses to the reproductive stage (R1) under ambient photoperiod and ∼27/22°C day/night temperature prior to drought treatment in a random block design. Following prior drought studies (Liu et al., 2015), drought stress was applied by withholding water consecutively for four weeks, whereas control plants were watered daily. Volumetric soil water content (SWC) was monitored regularly using a HydroSense II (Campbell Scientific, USA). Leaf and root tissues of treated and control plants (*n*=6 per group) were collected before the start of the treatment (week 0), after two weeks (week 2), and after four weeks (week 4) at a consistent time and immediately flash-frozen in liquid N_2_. To enable comparative data integration, samples for transcriptome and metabolite analyses originated from the same plant tissue samples, which were split for the different analyses.

### RNA isolation, transcriptome sequencing, and differential gene expression analysis

Total RNA was extracted from 100 mg of leaves or roots of Alamo and Cave-in-Rock plants (n=6) either drought-stressed or well-watered (control) using a Monarch^®^ Total RNA Miniprep Kit (New England Biolabs, USA) and subsequently treated with DNase I for genomic DNA removal. Following assessment of RNA integrity and quantitation using the Bioanalyzer 2100 RNA Nano 6000 Assay Kit (Agilent Technologies Inc., CA, USA), four of the six biological replicates with highest RNA quality were selected for sequencing. Preparation of cDNA libraries and transcriptome sequencing was performed at Novogene (Novogene Corporation Inc., USA). In brief, following RNA integrity analysis and quantitation, cDNA libraries were generated using a NEBNext^®^Ultra™RNA Library Prep Kit (New England Biolabs, USA) and sequenced on an Illumina Novaseq 6000 sequencing platform generating 40-80 million 150 bp paired-end reads per sample. Filtered, high-quality reads were aligned to the reference genome (*P. virgatum* var. Alamo AP13 v5.1) using HISAT2 (Kim et al. 2019a). Gene functional annotation was based on best matches to databases from Phytozome v13 (phytozome-next.jgi.doe.gov), including Arabidopsis, rice, Gene Ontology, and Panther, as well as in-house protein databases of biochemically verified terpene-metabolic enzymes (Pelot et al. 2018; Murphy and Zerbe 2020). Differentially expressed genes (DEGs) were identified based on padj < 0.05 and |log2 FC| > 1 as selection criteria. Statistical analyses were conducted in R and also plots and heatmaps were created using the ‘ggplot2’ and ‘pheatmap’ packages in R (cran-project.org, version 3.6.3).

### Metabolite extraction

Metabolite analysis followed previously established protocols (Li et al. (2020). Here, 100 mg tissue were ground to a fine powder in liquid N_2_ and metabolites were extracted with 1 ml 80% methanol containing 1 μM telmisartan internal standard by vortexing briefly and incubation for 16 h at 200 rpm and 4°C. Samples were centrifuged for 20 min (4000 g, 4°C) to remove solid particles and the supernatants transferred into fresh vials and stored at −80°C prior to LC-MS analysis.

### UPLC-ESI-QToF-MS analysis

Metabolite profiling was achieved by reversed-phase Ultra Performance Liquid Chromatography-Electrospray Ionization-Quadrupole Time-of-Flight mass spectrometry (UPLC-ESI-QToF-MS) analysis in positive and negative ionization mode on a Waters Acquity UPLC system equipped with a Waters Xevo G2-XS QToF MS (Waters, Milford, MA) and a Waters UPLC BEH C18 (1.7 μm x 2.1 mm x 150 mm) column. Chromatography was performed using 10 mM NH_4_HCO_2_ (in water; solvent A) and 100% acetonitrile (solvent B) as mobile phase and the following parameters: flow rate of 0.4 ml min^-1^; column temperature 40°C; 10 μl injection; method: 0-1 min (99% A/1% B), 1-15 min linear gradient to 1% A/99% B, 15-18 min (1% A/99%B), 18-20 min (99% A/1% B); QToF parameters: desolvation temperature of 350°C; desolvation gas flow rate at 600 L h^-1^; capillary voltage of 3 kV; cone voltage of 30 V. Mass spectra were acquired in continuum mode over *m*/*z* 50-1500 using data-independent acquisition (DIA) and MS^E^ with collision potential scanned at 20-80 V for the higher energy function.

The obtained DIA MS data were processed using Progenesis QI (V3.0, Waters) for quality control, chromatography alignment and mass feature extraction. Metabolite annotations were performed by matching *m*/*z* ratios, RT and fragmentation patterns against metabolite databases through Progenesis QI and LipidBlast (Kind et al. 2013). As a complimentary annotation approach, DDA (data dependent acquisition) was performed on a subset of the samples to obtain MS/MS spectral data for mainly the most abundant features. In addition, CANOPUS was used to predict chemical classes of the features based on their MS/MS information (Dührkop et al. 2021). Ion abundances of all detected features were normalized to the internal telmisartan standard based on five biological replicates. The normalized data (abundance > 300) were used for statistical analysis using MetaboAnalyst 5.0 (Pang et al. 2021).

### NMR analysis of terpenoid metabolites

About 200 g fresh root tissues of Cave-in-Rock plants were harvested. Compound purification was performed according to the method described in the Li et al (2020). The differences are the ethyl acetate and hexane phases (in which the diterpenoids were concentrated) were evaporated to dryness using a SpeedVac vacuum concentrator. The residue was re-dissolved in 8 mL of 95% methanol. Supernatants were transferred to LC vials. Purification was carried out as previously described using a C18 HPLC column (100 x 4,6 mm x 5µm). For NMR analysis, ∼0.4-0.8 mg of each HPLC purified compounds were dissolved in deuterated chloroform (CDCl_3_; Sigma-Aldrich, USA) containing 0.03% (v/v) tetramethylsilane (TMS). NMR 1D (^1^H and ^13^C) and 2D (HSQC, COSY, HMBC and NOESY) spectra were acquired as previously described (Pelot et al. 2018) on a Bruker Avance III 800 MHz spectrometer (Bruker Corporation, MA, USA) equipped with a 5 mm CPTCI cryoprobe using Bruker TopSpin 3.6.1 software and analyzed with MestReNova 14.1 software. Chemical shifts were calibrated against known TMS signals.

## Acknowledgements

We gratefully acknowledge Dr. Malay Saha at the Noble Research Institute for providing tillers for cultivation of Alamo AP13 and Cave-in-Rock, Dr. Daniel A. Jones at Michigan State University for his help with identification of diterpenoid features, and Dr. Andrew Muchlinski (Firmenich, San Diego, USA) for helpful discussions on the manuscript.

## Author Contributions

P.Z. and K.T. conceived the original research and oversaw data analysis; K.T. conducted plant drought stress experiments and transcriptome analysis; X.L. performed metabolite profiling and analysis; A.M., P.Y., and D.T. performed NMR structural analyses; D.D. and Y.C. assisted with plant harvesting, sampling, and sample processing; K.T. and P.Z. wrote the original article draft with editing by all authors. All authors have read and approved the manuscript.

## Data availability

The RNA-seq data were submitted to the Sequence Read Archive (SRA), accession no. PRJNA644234.

## Supporting Information

**Fig. S1:** Available water content (AWC) in the soil during the treatment

**Fig. S2:** Differentially expressed genes between all groups after four weeks of drought treatment

**Fig. S3:** Identification of significantly enriched metabolic pathways at the end of the treatment via GO term analysis

**Fig. S4:** LC-MS chromatograms and spectra of identified panicoloids

**Fig. S5:** Diterpenoid network in switchgrass

**Fig. S6:** NMR analysis of panicoloids A-C

**Table S1:** Complete list of differentially expressed genes (DEGs)

**Table S2:** Permutational multivariate analysis of variance (PERMANOVA) of a) gene expression levels and b) metabolite abundances

**Table S3:** Enrichment of GO terms and KEGG pathways in Alamo and Cave-in-Rock

**Table S4:** Complete list of the calculated FPKM (Fragments Per Kilobase of transcript per Million mapped reads) values for all genes

**Table S5:** List of all mass features from the positive mode LC-MS dataset that were selected for the downstream statistical analysis.

**Table S6:** Statistical analysis for detected LC-MS features of annotated specialized metabolites

