## Supplementary material for "Comparative transcriptomics and metabolomics reveal specialized metabolite drought stress responses in switchgrass (*Panicum virgatum* L.)": Fig. S1

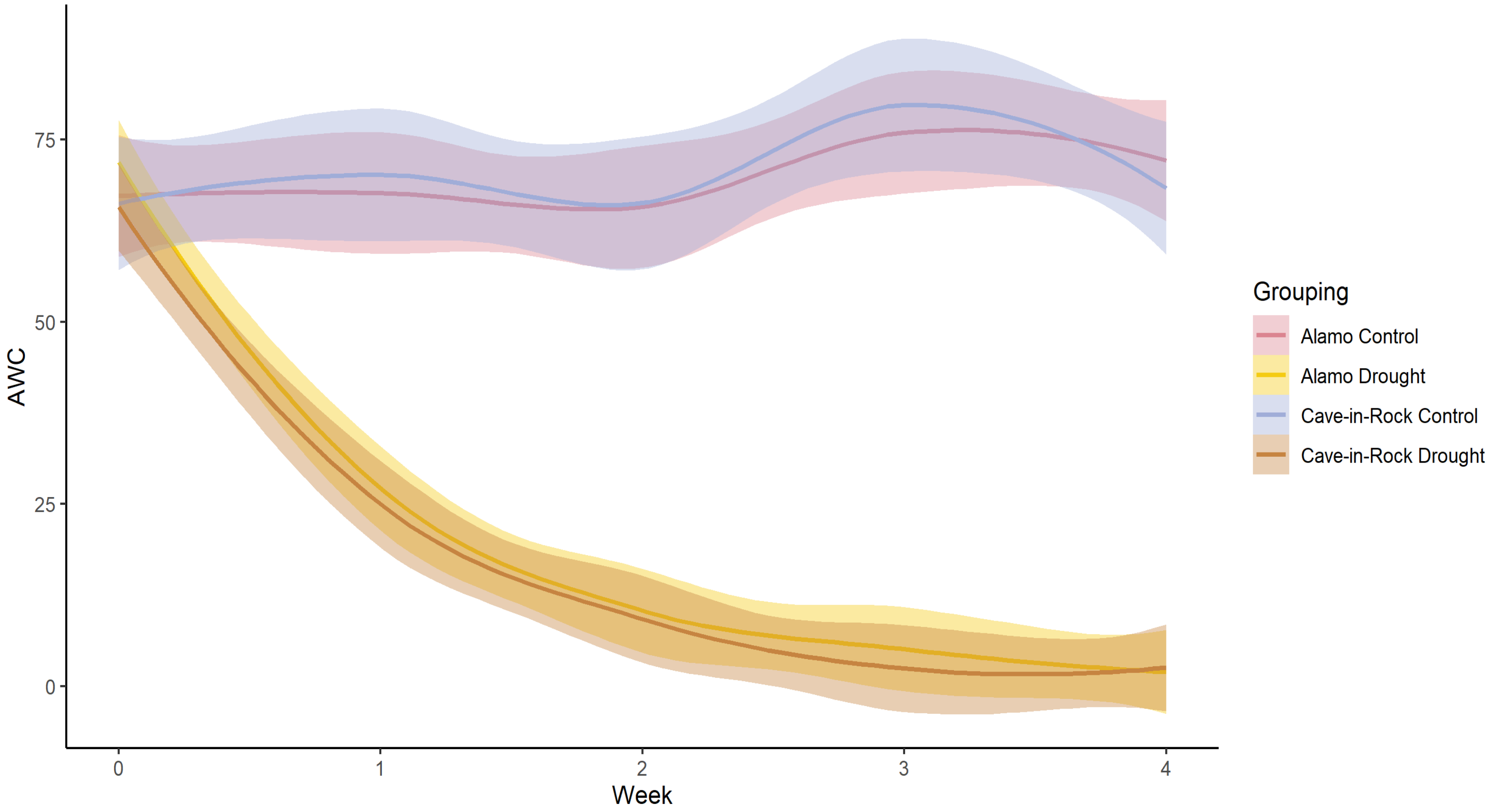


**Supporting Information Figure S1:** Available water content (AWC; in %) in the soil during the treatment.
