## Supplementary material for "Comparative transcriptomics and metabolomics reveal specialized metabolite drought stress responses in switchgrass (*Panicum virgatum* L.)": Fig. S2

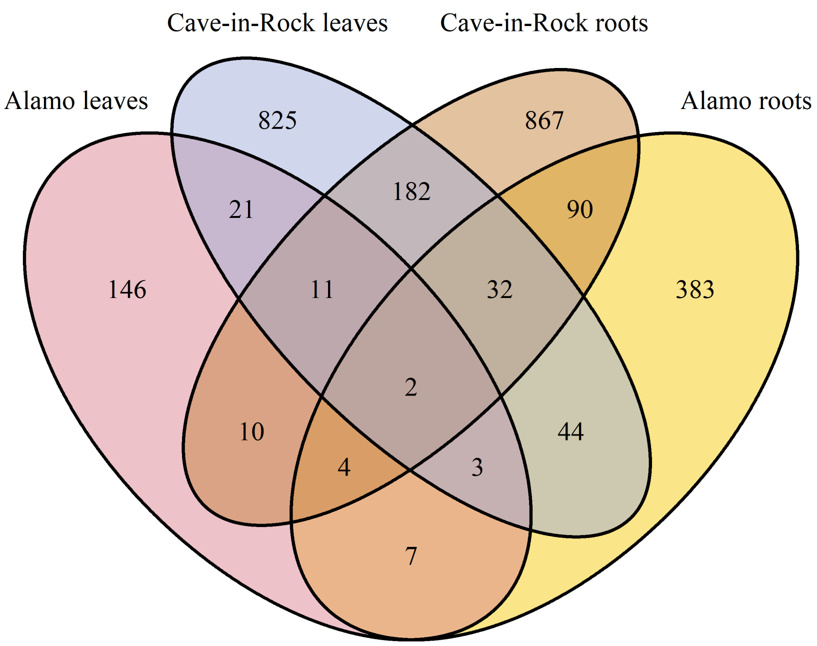


**Supporting Information Figure S2:** Number of differentially expressed genes (DEGs) between all groups after four weeks of drought treatment
