## Supplementary material for "Comparative transcriptomics and metabolomics reveal specialized metabolite drought stress responses in switchgrass (*Panicum virgatum* L.)": Fig. S3

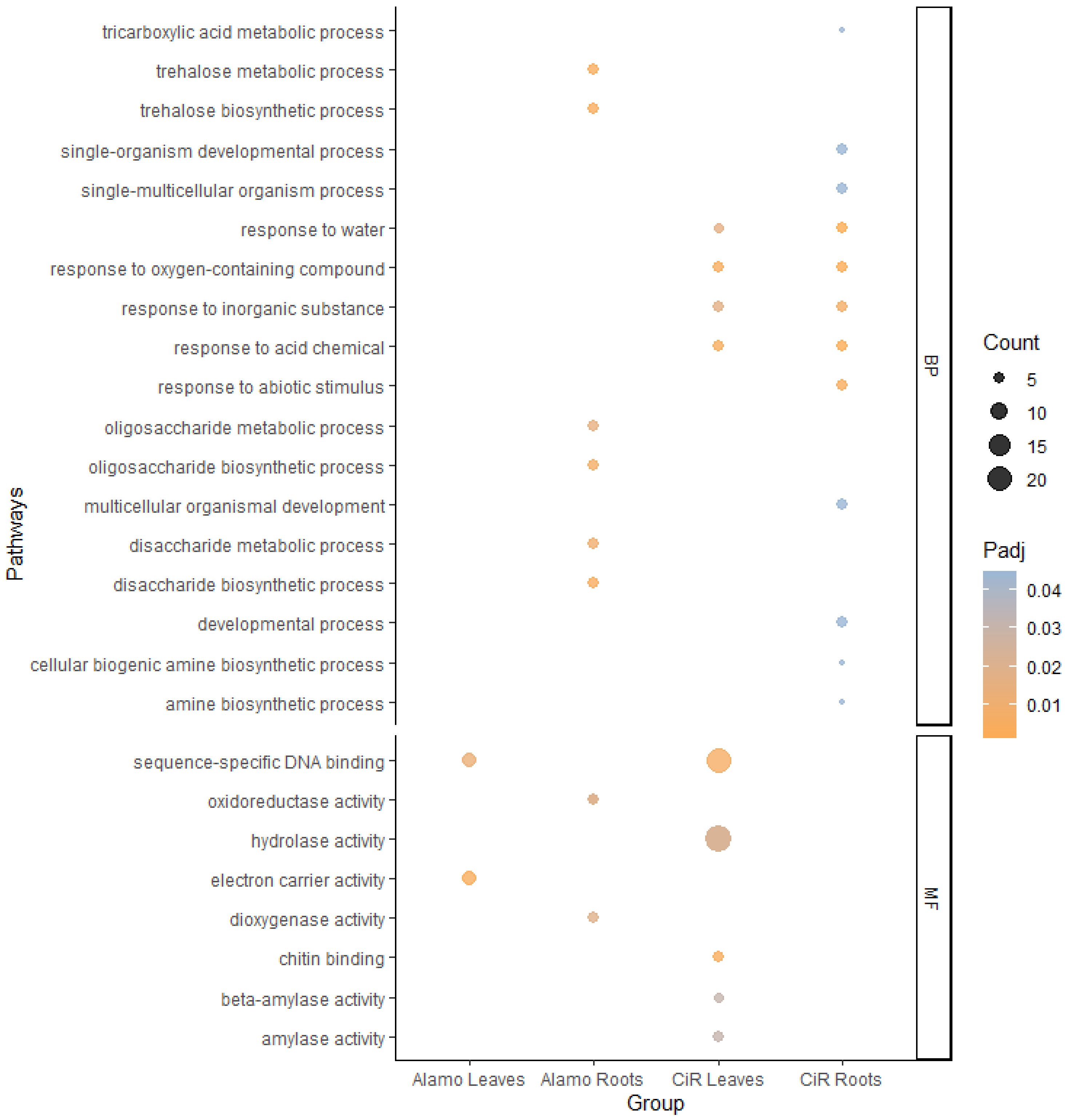


**Supporting Information Figure S3:** Identification of significantly enriched metabolic pathways at the end of the treatment via GO term analysis
