## Supplementary material for "Comparative transcriptomics and metabolomics reveal specialized metabolite drought stress responses in switchgrass (*Panicum virgatum* L.)": Fig. S4

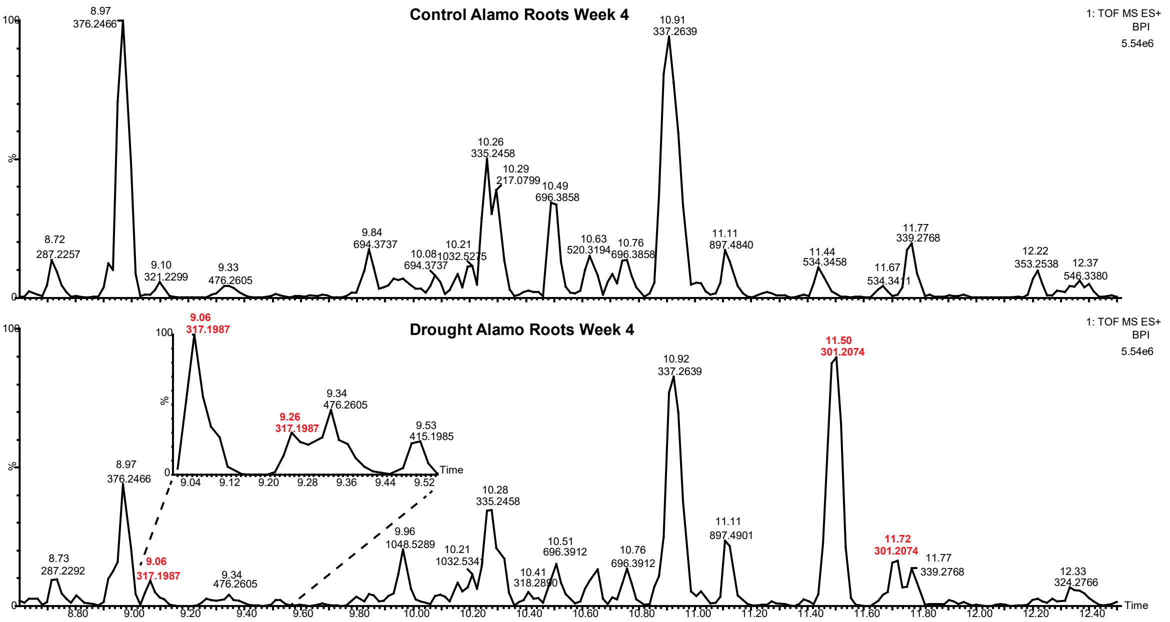
Control Alamo Roots Week 4
1: TOF MS ES+
BPI
5.54e6
8.97
376.2466
10.91
337.2639
10.26
335.2458
10.29
217.0799
10.49
696.3858
9.84
694.3737
8.72
287.2257
9.10
321.2299
9.33
476.2605
10.08
694.3737
10.21
1032.5275
10.63
520.3194
10.76
696.3858
11.11
897.4840
11.44
534.3458
11.67
534.3411
11.77
339.2768
12.22
353.2538
12.37
546.3380
Drought Alamo Roots Week 4
9.06
317.1987
9.26
317.1987
9.34
476.2605
9.53
415.1985
8.97
376.2466
8.73
287.2292
9.34
476.2605
9.96
1048.5289
10.21
1032.534
10.28
335.2458
10.41
318.2890
10.51
696.3912
10.76
696.3912
10.92
337.2639
11.11
897.4901
11.50
301.2074
11.72
301.2074
11.77
339.2768
12.33
324.2766
Time
%

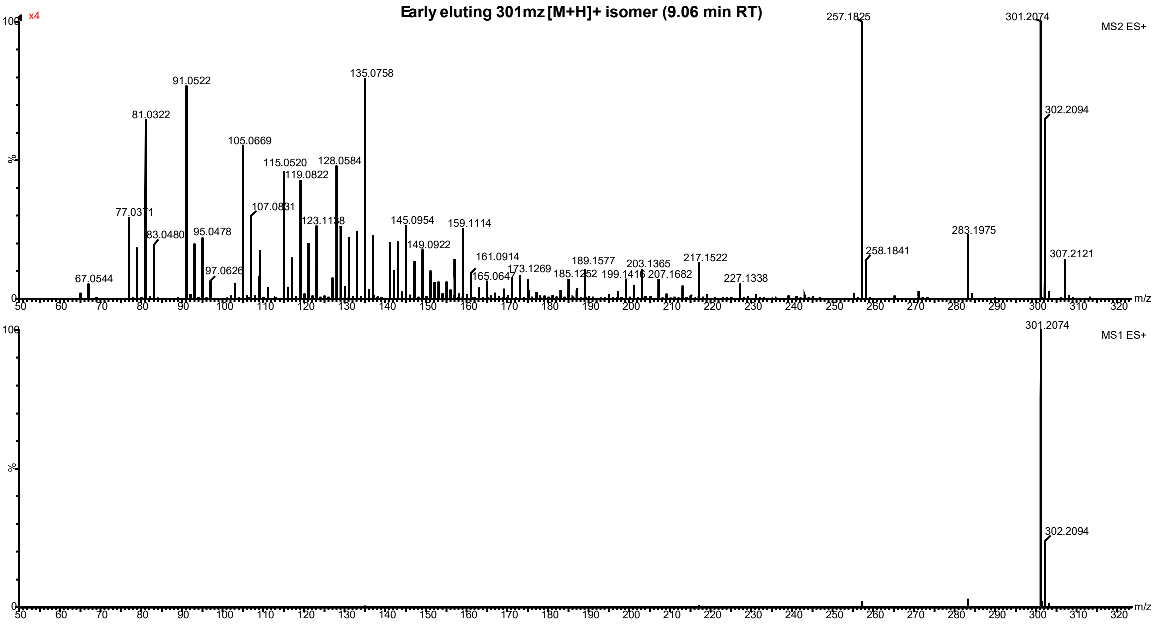
Early eluting 301mz [M+H]+ isomer (9.06 min RT)
x4
MS2 ES+
257.1825
301.2074
302.2094
135.0758
91.0522
81.0322
105.0669
115.0520
128.0584
119.0822
107.0831
77.0371
83.0480
95.0478
123.1138
145.0954
159.1114
149.0922
97.0626
67.0544
161.0914
165.064
173.1269
185.1252
189.1577
199.1416
203.1365
207.1682
217.1522
227.1338
258.1841
283.1975
307.2121
MS1 ES+
301.2074
302.2094
m/z
%

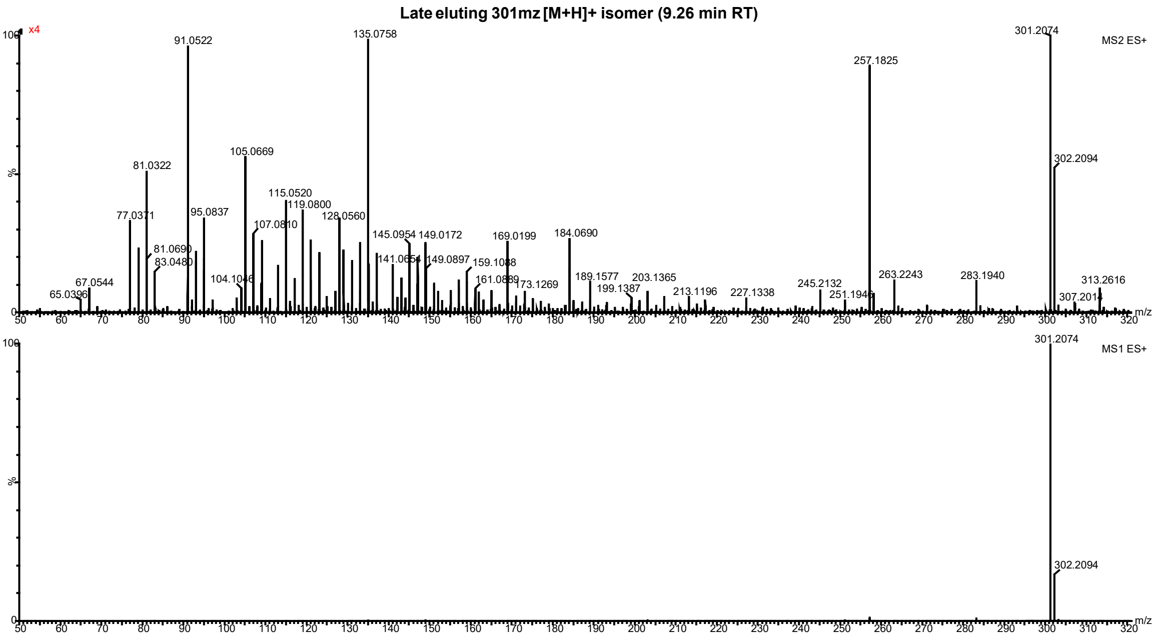
Late eluting 301mz [M+H]+ isomer (9.26 min RT)
x4
MS2 ES+
91.0522
135.0758
301.2074
257.1825
302.2094
105.0669
81.0322
115.0520
119.0800
77.0371
95.0837
107.0810
128.0560
145.0954
149.0172
169.0199
184.0690
81.0690
83.0480
141.0654
149.0897
159.1048
161.0889
173.1269
189.1577
199.1387
203.1365
213.1196
227.1338
245.2132
251.1946
263.2243
283.1940
313.2616
307.2014
104.1046
67.0544
65.0396
MS1 ES+
301.2074
302.2094
m/z
%

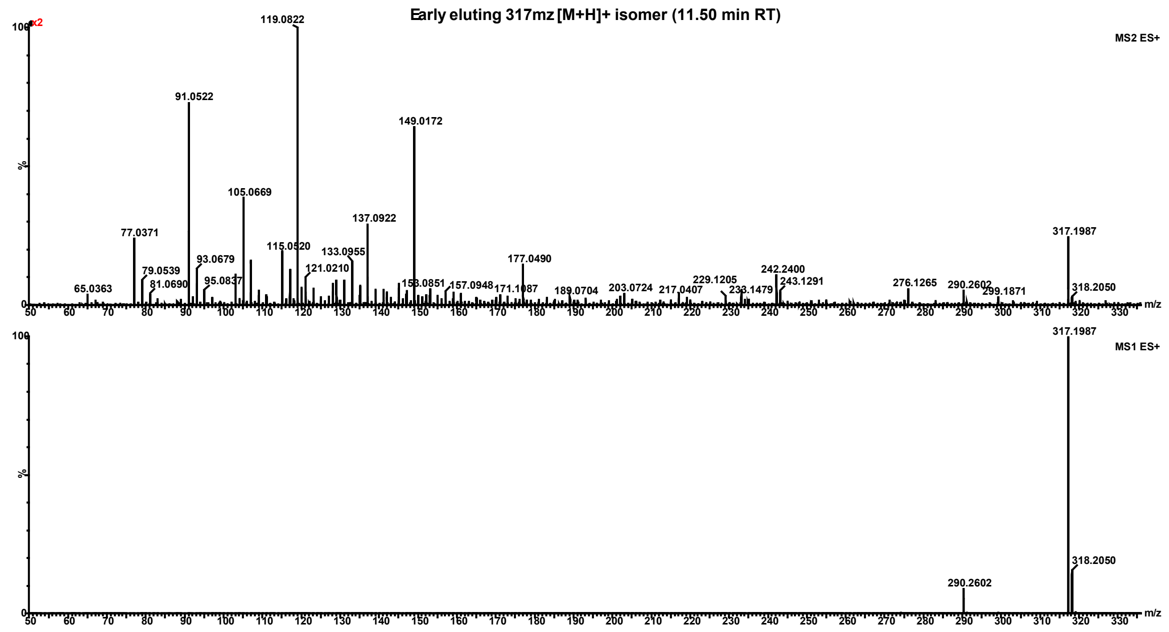


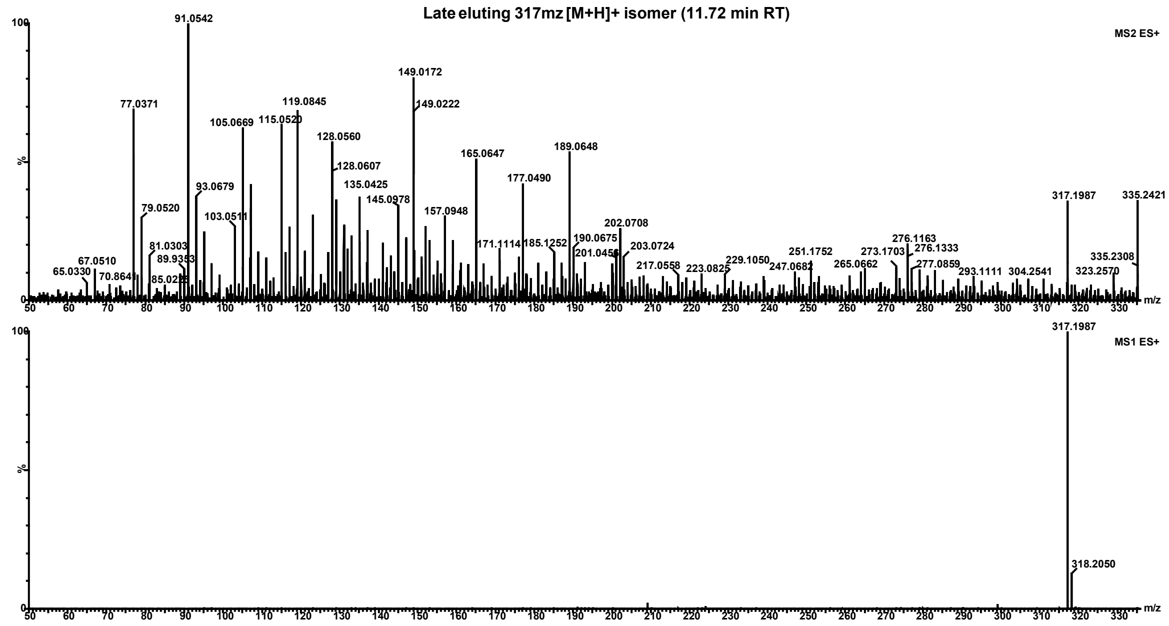


**Fig. S4: LC-MS chromatograms and spectra of identified panicoloids.** Shown chromatograms are example chromatograms from Alamo roots after four weeks of either well-watered (control) or drought-stressed conditions (drought) with panicoloid features highlighted in red. Chromatograms and spectra were obtained via LC-MS ESI analysis of compounds isolated from switchgrass root tissue.
