## Supplementary material for "Comparative transcriptomics and metabolomics reveal specialized metabolite drought stress responses in switchgrass (*Panicum virgatum* L.)": Fig. S5

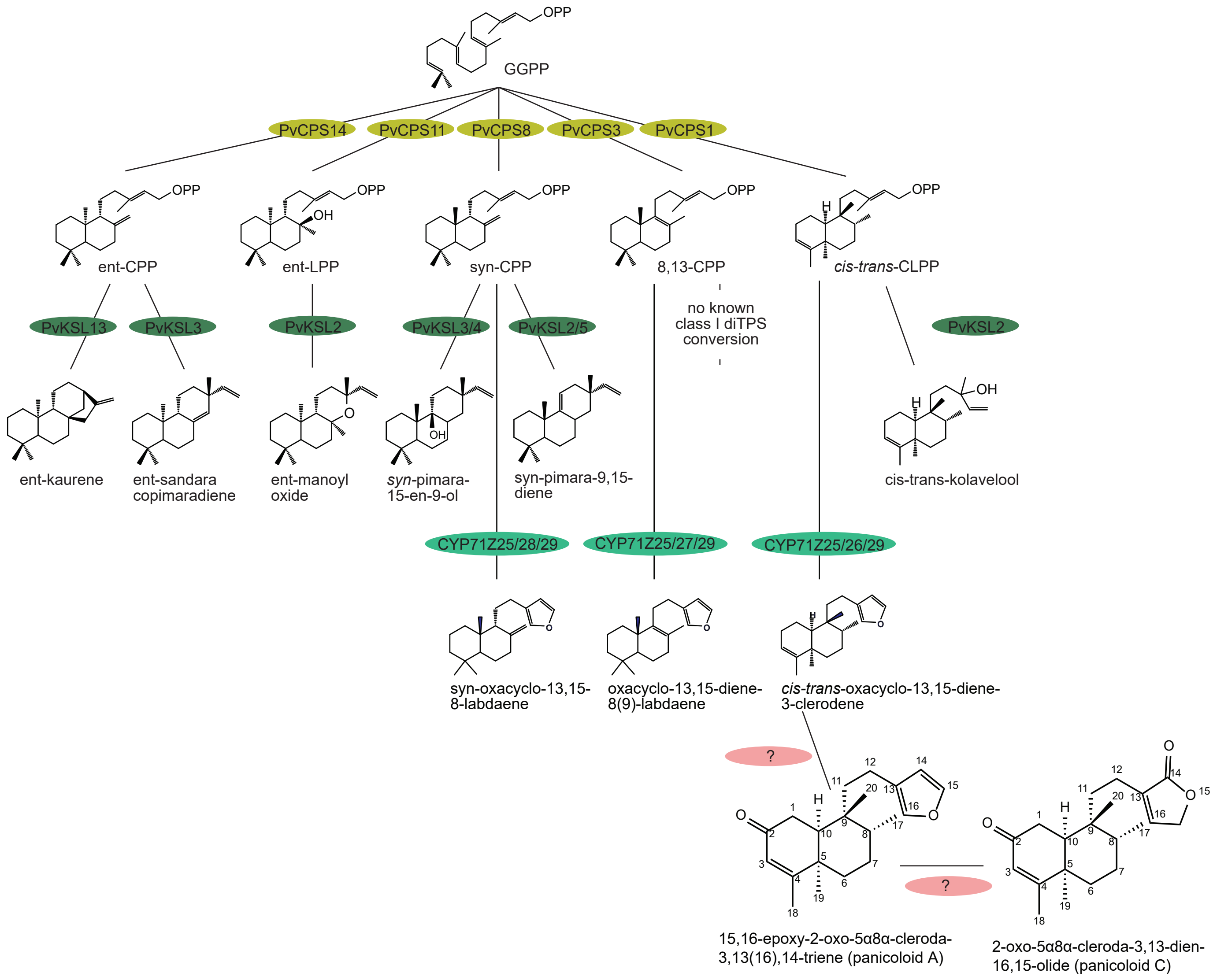


**Supporting Information Figure S5:** So far identified diterpenoid network in switchgrass. Abbreviations: GGPP, geranylgeranyl pyrophosphate; CPP, copalyl pyrophosphate; LPP, labdadienyl pyrophosphate; CLPP, clerodienyl pyrophosphate; CPS, copalyl diphosphate synthase; KSL, kaurene synthase-like.
