## Supplementary material for "Comparative transcriptomics and metabolomics reveal specialized metabolite drought stress responses in switchgrass (*Panicum virgatum* L.)": Fig. S6

**Supporting Information Fig. 6: NMR analysis of panicoloids A-C**

**Panicoloid A** - $C_{20}H_{28}O_2$ - diterpenoid 301 m/z $^1$H and $^{13}$C NMR (800 MHz, Chloroform-*d*)

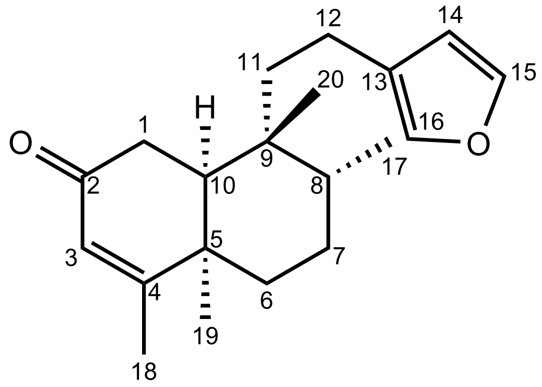

| Position | δC (ppm) | δH (ppm) | J (Hz) |
|---|---|---|---|
| 1 a<br>b | 35.5 | 2.60 (m)<br>2.73 (dd) | 18.3, 6.6 |
| 2 | 199.2 | | |
| 3 | 128.5 | 5.88 (s) | |
| 4 | 168.9 | | |
| 5 | 39.4 | | |
| 6 a<br>b | 29.8 | 1.52 (m)<br>1.86 (dt) | 14.5, 3.7 |
| 7 a<br>b | 26.5 | 1.34 (dt)<br>1.64 (tt) | 13.5, 3.7<br>13.8, 3.6 |
| 8 | 35.1 | 1.71 (tq) | 7.4, 3.7 |
| 9 | 38.9 | | |
| 10 | 46.7 | 1.94 (dd) | 6.7, 2.2 |
| 11 a<br>b | 38.9 | 1.30 (ddd)<br>1.78 (m) | 13.4, 10.2, 6.7 |
| 12 | 18.1 | 2.33 (m) | |
| 13 | 125.4 | | |
| 14 | 110.9 | 6.26 (dd) | 1.9, 0.9 |
| 15 | 142.8 | 7.34 (t) | 1.7 |
| 16 | 138.5 | 7.20 (dq) | 1.9, 1.0 |
| 17 | 14.5 | 1.00 (d) | 7.1 |
| 18 | 20.9 | 1.97 (d) | 1.4 |
| 19 | 31.07 | 1.26 (s) | |
| 20 | 22.8 | 0.90 (s) | |

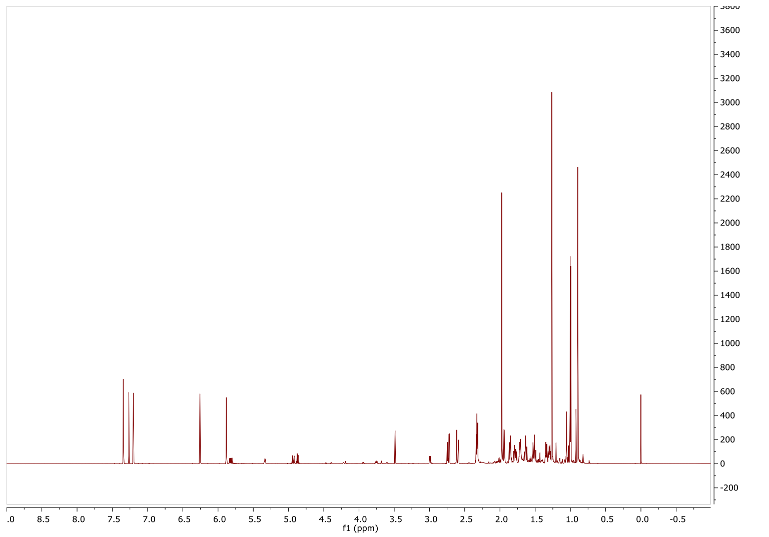
f1 (ppm)

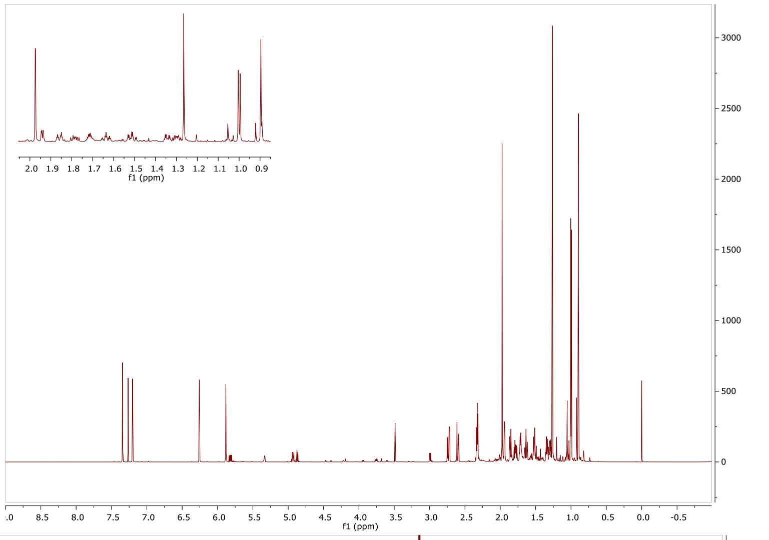
f1 (ppm)
f1 (ppm)

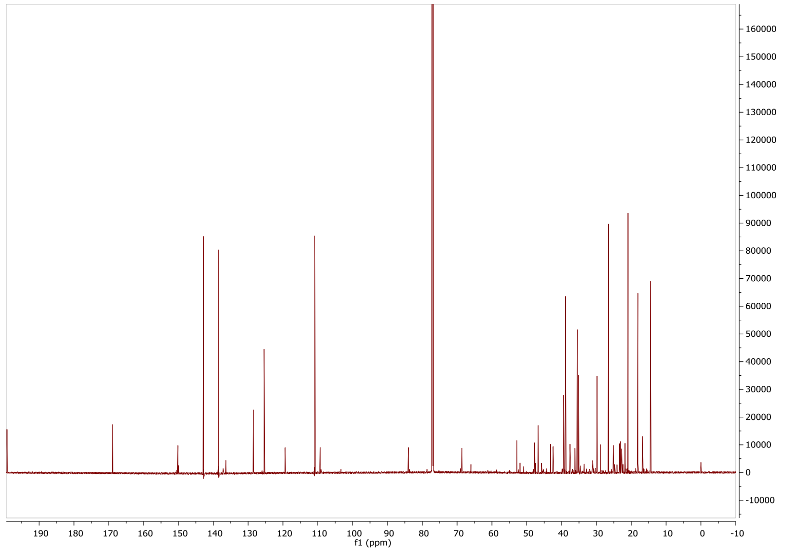
f1 (ppm)

**Panicoloid B** - $C_{20}H_{28}O_2$ - diterpenoid 301 m/z $^1$H and $^{13}$C NMR (800 MHz, Chloroform-*d*)

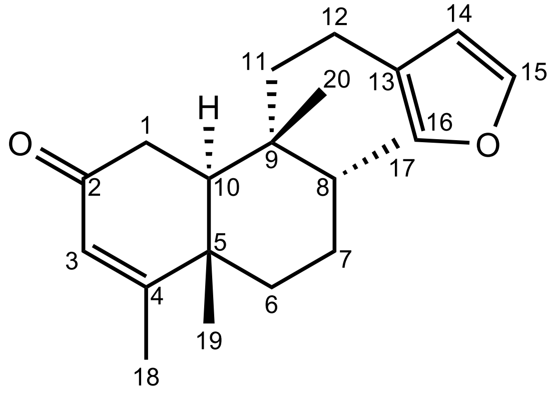

| Position | δC (ppm) | δH (ppm) | J (Hz) |
|---|---|---|---|
| 1 a<br>b | 34.8 | 2.34 (m)<br>2.44 (dd) | <br>17.6, 3.6 |
| 2 | 200.4 | | |
| 3 | 125.5 | 5.71 (t) | 1.2 |
| 4 | 172.9 | | |
| 5 | 40.0 | | |
| 6 a<br>b | 29.0 | 1.56 (dt)<br>1.61 (m) | 13.2, 3.6 |
| 7 | 25.0 | 1.43 (dq)<br>2.04 (tt) | 14.8, 3.4<br>14.0, 4.1 |
| 8 | 34.4 | 1.76 (m) | |
| 9 | 37.7 | | |
| 10 | 44.5 | 1.99 (dd) | 14.0, 3.6 |
| 11 a<br>b | 39.4 | 1.27 (m)<br>1.61 (m) | |
| 12 | 18.0 | 2.37 (m) | |
| 13 | 125.3 | | |
| 14 | 110.8 | 6.24 (dd) | 1.8, 0.9 |
| 15 | 142.8 | 7.34 (t) | 1.7 |
| 16 | 138.4 | 7.19 (dq) | 1.9, 1.0 |
| 17 | 14.4 | 0.98 (d) | 7.1 |
| 18 | 18.9 | 1.90 (d) | 1.4 |
| 19 | 18.9 | 1.17 (s) | |
| 20 | 20.0 | 1.07 (s) | |

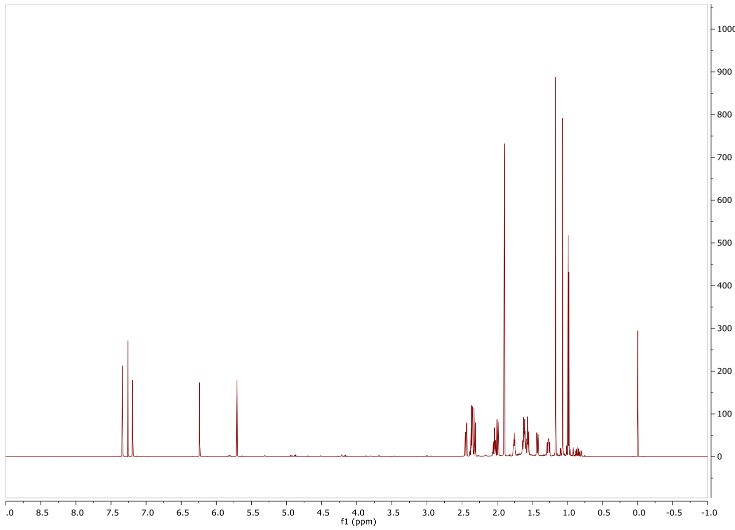
1000
900
800
700
600
500
400
300
200
100
0
.0 8.5 8.0 7.5 7.0 6.5 6.0 5.5 5.0 4.5 4.0 3.5 3.0 2.5 2.0 1.5 1.0 0.5 0.0 -0.5 -1.0
f1 (ppm)

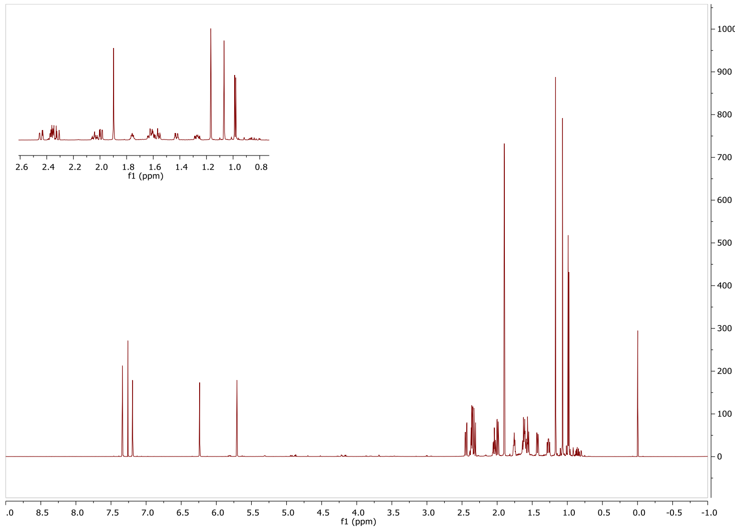
2.6 2.4 2.2 2.0 1.8 1.6 1.4 1.2 1.0 0.8
f1 (ppm)
1000
900
800
700
600
500
400
300
200
100
0
.0 8.5 8.0 7.5 7.0 6.5 6.0 5.5 5.0 4.5 4.0 3.5 3.0 2.5 2.0 1.5 1.0 0.5 0.0 -0.5 -1.0
f1 (ppm)

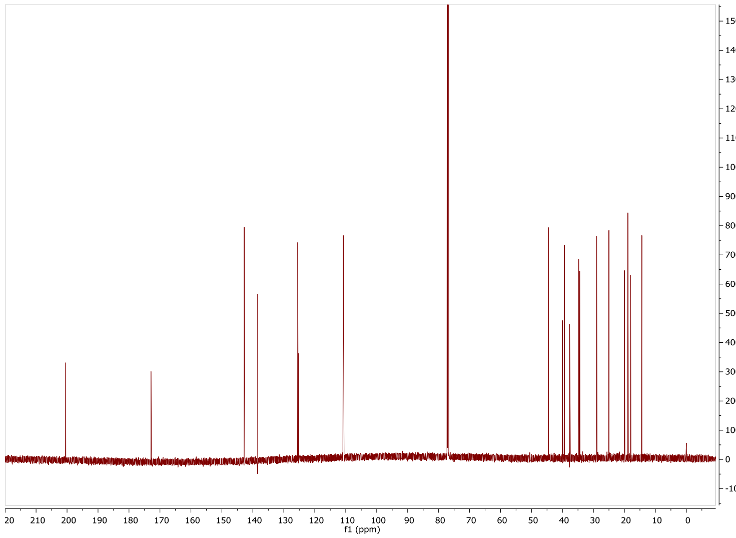
15000
14000
13000
12000
11000
10000
9000
8000
7000
6000
5000
4000
3000
2000
1000
0
-1000
20 210 200 190 180 170 160 150 140 130 120 110 100 90 80 70 60 50 40 30 20 10 0
f1 (ppm)

**Panicoloid C** – diterpenoid 317 m/z_1 $^{1}$H and $^{13}$C NMR (800 MHz, Chloroform-*d*)

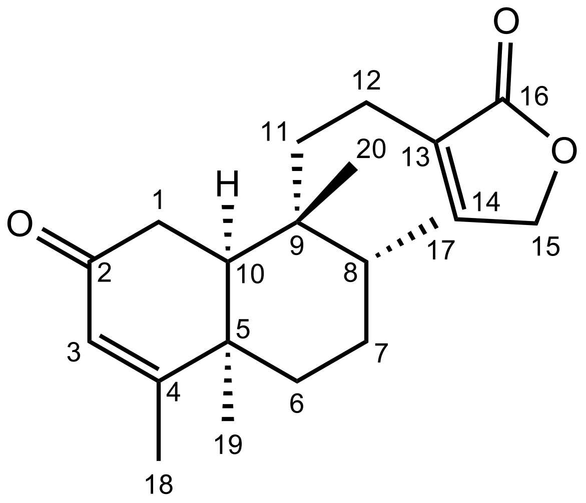

| Position | $\delta_C$ (ppm) | $\delta_H$ (ppm) | J (Hz) |
|---|---|---|---|
| 1 a<br>b | 35.4 | 2.56 (m)<br>2.76 (dd) | 18.3, 6.7 |
| 2 | 199 | | |
| 3 | 128.5 | 5.88 (s) | |
| 4 | 169 | | |
| 5 | 39.4 | | |
| 6 a<br>b | 29.7 | 1.51 (m)<br>1.86 (dt) | 14.5, 3.5 |
| 7 a<br>b | 26.5 | 1.33 (m)<br>1.64 (m) | |
| 8 | 35.1 | 1.70 (m) | |
| 9 | 38.8 | | |
| 10 | 46.7 | 1.96 (m) | |
| 11 a<br>b | 36.0 | 1.36 (m)<br>1.70 (dd) | 12.7, 4.5 |
| 12 | 18.9 | 2.21 (m) | |
| 13 | 134.9 | | |
| 14 | 143.8 | 7.09 (m) | |
| 15 | 70.2 | 4.78 (q) | 2.1 |
| 16 | 174.4 | | |
| 17 | 14.3 | 1.01 (d) | 7.1 |
| 18 | 21.0 | 1.97 (m) | |
| 19 | 31.1 | 1.27 (s) | |
| 20 | 22.7 | 0.88 (s) | |

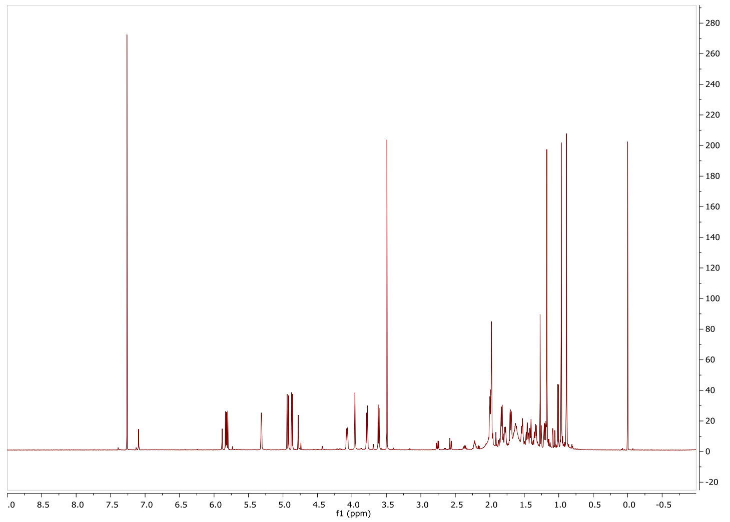
f1 (ppm)

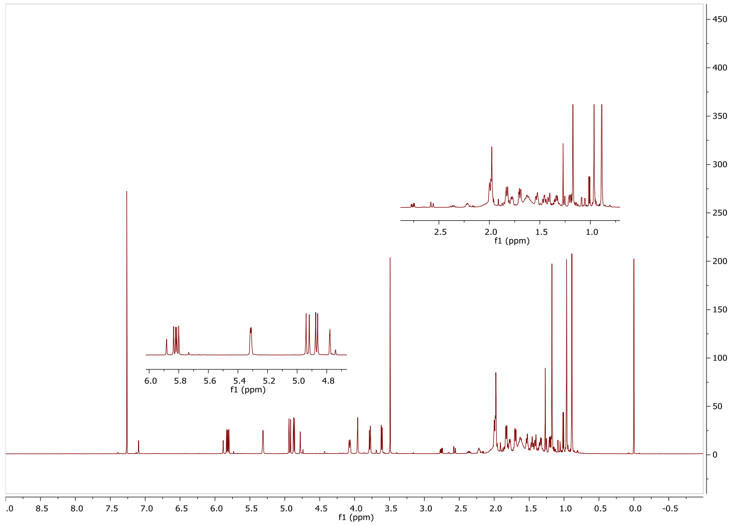
f1 (ppm)
f1 (ppm)
f1 (ppm)

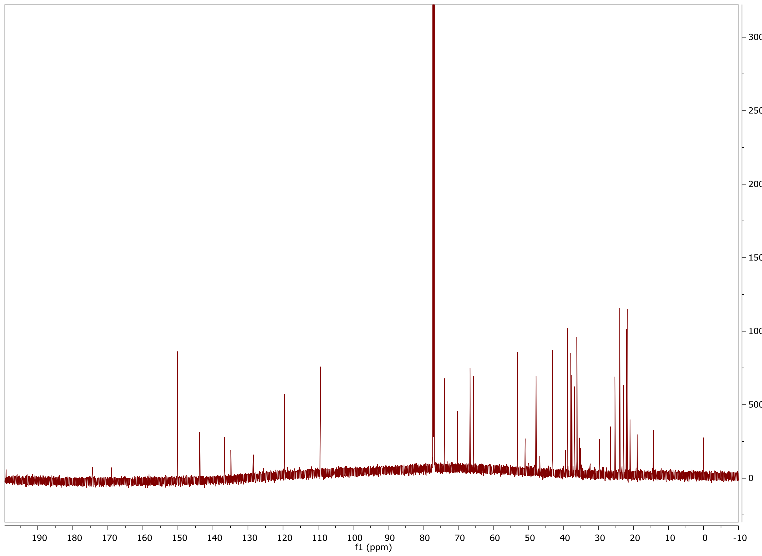
f1 (ppm)

301 m/z (Top) and 317 m/z_1 (Bottom) $^{1}H$ NMR (800 MHz, Chloroform-*d*)

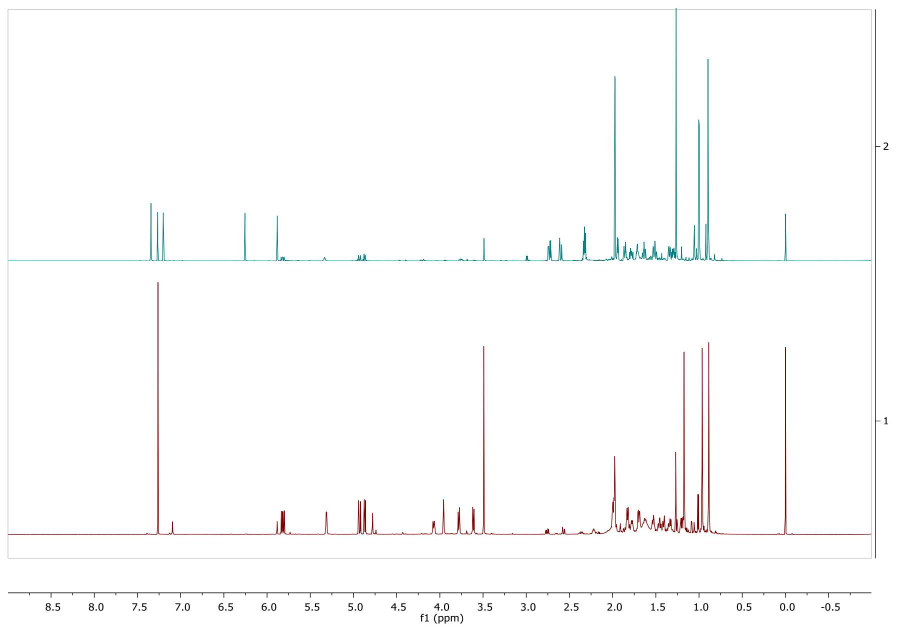

301 m/z (Top) and 317 m/z_1 (Bottom) $^{13}C$ NMR (800 MHz, Chloroform-*d*)

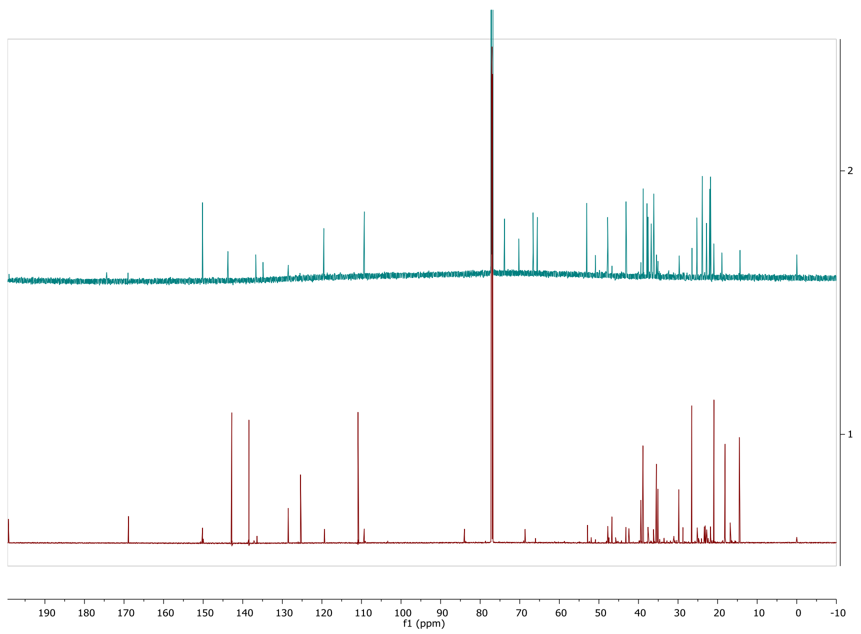
